# Minimal Computational Framework for Systematic Identification of Antimicrobial Targets

**DOI:** 10.64898/2026.05.24.727537

**Authors:** Sergio A. Hassan

## Abstract

Systematic identification of antimicrobial targets remains a major challenge, as discovery still relies largely on empirical, resource-intensive approaches with limited efficiency. We present a method for identifying antimicrobial targets based on protein dynamics, enabling rational polypharmacology. The approach spans multiple biological scales, from taxa (genus and species) to biological networks, including network hubs and edges, their constituent proteins, protein binding sites, and their conformational states. It is grounded in the premise that coordinated intervention across multiple, optimally selected targets, using combinations of compounds at safe or submaximal doses, can achieve therapeutic effects while reducing toxicity and limiting mutational escape. A survey of known antimicrobials indicates that a small number of recurrent protein-level mechanisms account for most disruptions of microbial survival. We introduce metrics to detect these mechanisms across a pathogen proteome and describe a streamlined, modular workflow for target identification and prioritization that is optimized for ease of deployment and naturally interfaces with downstream applications such as molecular screening and *de novo* design.

## Introduction

Current antimicrobials target only a small fraction of a pathogen’s proteome, leaving many biological networks essential for survival unexplored. Moreover, all clinically used antimicrobials have been discovered experimentally, with their parent compounds typically identified by chance. Rational design remains limited by gaps in structural information and incomplete knowledge of effective targets. High-throughput screening of chemical libraries and soil-derived microorganisms offers a more systematic approach to discovering new chemical classes, but these methods remain inefficient: chemical libraries cover only a tiny fraction of chemical space, and most microbial diversity in soil, the main source of natural products, cannot be cultured with current techniques.^1^ Advances in cultivation methods could accelerate progress, as illustrated by the recent discovery of peptides with strong antimicrobial activity.^2^ Despite these successes, the inefficiency of traditional approaches has contributed to the antibiotic pipeline crisis, a problem further exacerbated by microbial evolution and the emergence of resistant strains.^3,4^

Computational approaches complement experimental methods by leveraging large virtual libraries of molecules. Although high-throughput virtual screening has not yet yielded new chemical classes of antimicrobials, it can identify promising lead compounds for early-stage drug discovery and for optimizing existing agents. However, many of these leads fail during subsequent development because activity predicted in silico or observed in vitro often does not translate to the physiological complexity of in vivo systems,^5^ resulting in toxicity, off-target effects, unfavorable pharmacokinetic and pharmacodynamic profiles, and additional adverse outcomes. Rapid development of resistance further reduces their effectiveness.^6,7^ Expanding virtual libraries, now reaching billions to trillions of molecules,^8,9^ is important but may not be sufficient, because longstanding challenges, often justified in practice by the need for computational efficiency, are likely to persist. These include limitations in binding-site modeling, insufficient conformational sampling, and simplified scoring functions, all of which can lead to false positives or, more problematically, false negatives that may prematurely exclude viable lead compounds. Data-driven computational approaches, particularly those based on machine learning and artificial intelligence,^10,11^ are currently being developed as promising alternatives in drug discovery. Their practical impact remains an active area of evaluation, in part due to the relative scarcity of high-quality, experimentally validated data available for model training. To place this challenge in perspective, since the advent of modern antimicrobial therapy in the 1940s, the total number of approved agents remains limited, on the order of 150 antibacterials, 100 antivirals, 30 antiparasitics, and 20 antifungals. These estimates refer to unique active compounds and generally include agents approved at any point in time, regardless of whether they were later withdrawn, superseded, or rendered less effective due to the emergence of resistance. Even fewer chemical classes exist; for example, current antifungal agents are commonly grouped into four major classes (polyenes, azoles, echinocandins, and allylamines), underscoring the limited scaffold diversity and constrained chemical space that data-driven approaches must navigate.

Overall, the likelihood of discovering an effective antimicrobial remains low, whether through experimental or computational approaches (see Discussion). There is growing consensus that progress requires novel strategies^12–16^ and options are being explored.^2,10,11,17,18^ The computational framework proposed here addresses this challenge by shifting the focus from identifying lead compounds to identifying multiple potential targets. The design or use of therapeutic agents that act on multiple targets or disease pathways is referred to as polypharmacology.^19–21^ This represents a paradigm shift from the traditional ‘one drug, one target’ model to strategies in which single agents or combinations of agents act on multiple targets (see Discussion).

Throughout this paper, ‘target’ may refer to entities across multiple biological scales, including a genus or species, biological networks, subnetworks or pathways, network hubs or edges, constituent proteins, protein binding sites, or the conformational substates thereof. Membranes and lipid molecules, important targets for both established and emerging antimicrobials (e.g., polyenes and membrane-disrupting peptides,^22^ are excluded here because they require distinct methodological approaches.^23^ The core of the method is the systematic scanning of a given target and the extraction of metrics representing known mechanisms of action. The Application section of the paper illustrates the use of the method in practice, validates its ability to recover known antimicrobial targets, and demonstrates its potential to identify and prioritize putative targets.

A minimal, self-contained pipeline, built from free, open-source components and integrated with public databases, is provided to enable systematic exploration of microbial proteomes. Designed for simplicity, ease of deployment, and adaptability to both institutional clusters and distributed resources, the pipeline is optimized for high-performance and high-throughput computing workflows. Its modular architecture and core algorithmic logic are intended to ensure long-term robustness and reduce susceptibility to technological obsolescence, even in the context of fast-evolving computational tools and methods.

## I. Method

The development of the method was guided by both conceptual and technical considerations, with the former discussed in detail in the Discussion. The method uses protein conformational dynamics to derive a set of metrics capturing mechanistic behaviors exploited by known antimicrobials. It identifies vulnerable regions in proteins or complexes where perturbation is likely to disrupt the biological networks in which they operate. Network function and, in some cases, topology can be modulated by targeting specific vertices, edges, or hubs with one or more compounds.^24–28^ When concerted attack is optimized and sustained, it can, depending on the network’s robustness, either disable the network entirely or significantly impair its function. Both outcomes are desirable for pathogen control (see Discussion). Identifying multiple targets increases the likelihood of early-stage detection of antimicrobials suitable for multidrug combination strategies, ^19–21^ while lowering the risk of host toxicity and the development of pathogen resistance mechanisms (see Discussion).

The workflow is summarized in Fig. 1A, with a detailed flowchart provided in the Supporting Information. References, versions, and URLs for all software packages, simulation engines, and databases used in this work are provided in the Supporting Information, along with the DOI of the repository containing a prototype implementation of the pipeline. It begins with input data supplied by the user or retrieved from public databases and ends with the generation of three types of output. Workflow access is managed by submitting a `<session>` file (e.g., a plain-text `session.str`) containing any number of requests, one per line. Conceptually, each request follows the form shown in **Scheme 1** (see Applications and SI for details). A biological unit is first selected as the query (QUERy), and optionally, all proteins physically interacting with each protein contained in the query can be included (INTEractors). Biological units can range from a single protein sequence or a metabolic pathway up to an entire organism. For each protein collected, a series of metrics, selected independently for the queried proteins and their interactors, is computed to identify potential intervention targets. These metrics, described below, include physicochemical features derived from molecular dynamics simulations, ranging from coarse properties such as surface topography to subtle atomistic details, including internal water networks capable of shuttling protons, each chosen to capture a functional aspect of the protein’s behavior.

**Figure 1.**
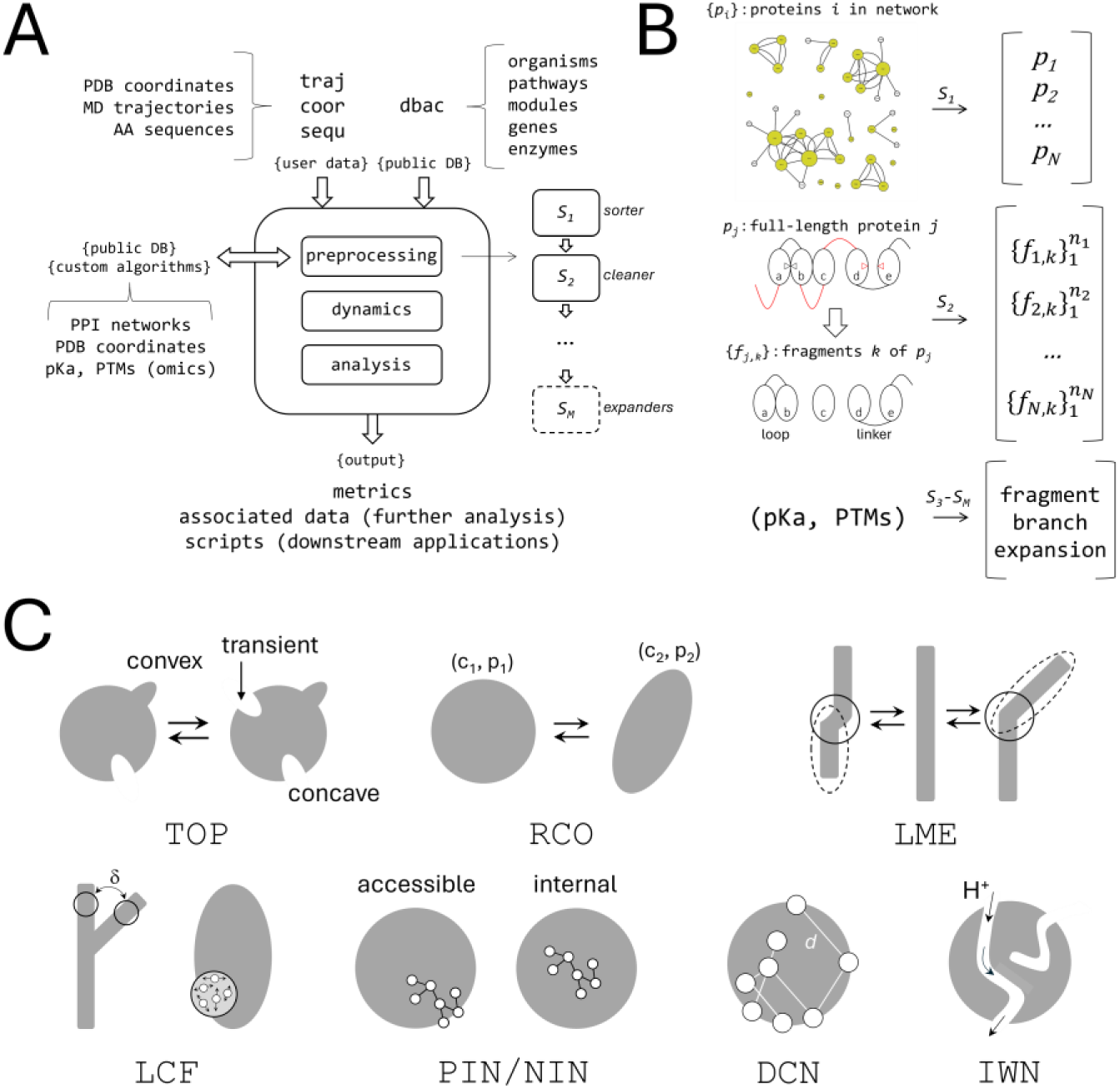
(A) Overview of the workflow (see detailed flowchart in SI). Inputs include user-provided dynamical trajectories, model structures, protein sequences, and database queries across multiple biological levels. Outputs comprise quantitative metrics and associated data for further analysis, along with scripts for visualization and downstream calculations (see flowchart in SI). (B) Preprocessing modules: the upper panel shows a protein–protein interaction (PPI) network with proteins {*p_j_*} ranked by node degree (default) or by other centrality measures; double edges indicate proteins already in the query, highlighting sites of particular interest (see III.2). The middle panel illustrates structure sanitization of predicted full-length models, in which low-confidence segments (red) are removed based on length and inter-domain alignment criteria (red triangles), producing *n_j_* fragments {*f_j,k_*} from each protein *p_j_*. The lower panel depicts optional combinatorial fragmentation based on PTM, pKa shifts, or cis–trans isomerization (see Discussion). (C) Minimal metrics from molecular dynamics simulations: (i) surface topography (`TOP`) identifies pockets and crevices, including transient features; (ii) representative conformations and occupancies (`RCO`) capture dominant conformers and their prevalence; (iii) local mobile elements (`LME`) detect hinges, proximity changes (not shown), and backbone distortions (solid circles) while ignoring global motions (dashed); (iv) local conformational fluctuations (`LCF`) quantify side-chain flexibility, treating the local patch as fluid while ignoring global fluctuations (δ); (v) polar (`PIN`) and nonpolar (`NIN`) interaction networks highlight connectivity patterns, with surface-extending interactions potentially vulnerable to perturbation; (vi) distance correlation networks (`DCN`) reveal potential long-range communication pathways; and (vii) internal water networks (`IWN`) identify buried water chains that may act as proton wires. All metrics are mapped as heatmaps onto a common reference structure (see Applications and SI); the rationale for selecting these metrics is provided in the Discussion section.

**Scheme 1:**
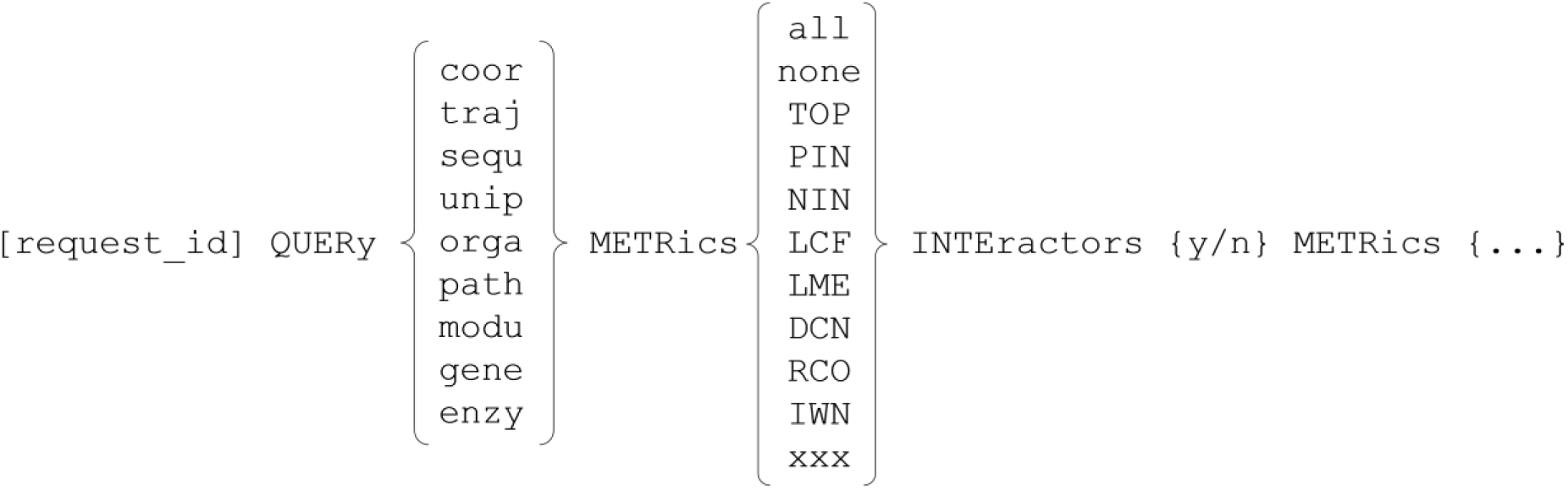
Conceptual structure of a `<session>` file request. All results are stored in dedicated directory, `request.[request_id]`/

Each request falls into one of four types (Fig. 1A): (1) coordinates of a model (`coor`), (2) trajectory from a dynamics simulation (`traj`), (3) amino acid sequence (`sequ`), or (4) accession code from a public database (`dbac`). The `coor`, `sequ`, and `dbac` types ultimately produce trajectories that feed into the downstream flow of `traj`. The model specified in `coor` uses PDB coordinates, whereas the sequence in `sequ` is provided in FASTA format. The trajectory queried via `traj` may contain proteins, peptides, membranes, ligands, ions, cosolutes, and other molecules; however, only the amino acid–based components, including proteins or peptides, whether linear or polycyclic, are analyzed downstream. By default, only the 20 naturally occurring amino acids are included; however, unnatural, unusual, or modified amino acids can be accommodated through simple modifications of a few scripts. Two databases are supported for `dbac`: UniProt (`unip`) and KEGG (`kegg`); see **Scheme 1** and *Session requests*. In `unip`, accession codes (AC) specify individual proteins, whereas `kegg` identifiers (ID) specify organisms (`orga`), pathways (`path`), modules (`modu`), genes (`gene`), or enzymes (`enzy`), covering most relevant biological entities while minimizing redundancy. The pipeline’s modular design enables integration of additional databases via API requests (see flowchart in SI).

For `dbac`, an optional parameter (`INTE`) queries the STRING database to retrieve interactors of the queried proteins. When multiple proteins are queried (e.g., via `path`), interactors of all proteins in the query are included. By default, only proteins experimentally confirmed with high confidence (score ≥ 0.75) to belong to the same physical complex are retained. This strict PPI network definition improves protocol efficiency by focusing on interactions most likely to influence network function. All queried proteins and their interactors (if `INTE` is enabled) then proceed to the coordinate-generation stage. Proteins are assigned UniProt ACs, and the pipeline first searches the AlphaFold2 database for a model. If a model is available, the coordinates and predicted errors of the top-ranked model are retrieved. Otherwise, a local AF2 implementation can be invoked through `sequ`. This type of query is particularly relevant for viruses, which are not represented in the database, and for many pathogenic organisms, as only the most common microbes are included (see Applications).

Before the simulation stage, all proteins from `dbac` undergo preprocessing. In the first module (*sorter*, *S_1_*), proteins are ranked by their importance within the PPI network, and a script is generated to visualize the network (Fig. 1B; upper panel). Hub nodes, i.e., highly interconnected proteins, are prioritized. Queried proteins are generally ranked above interactors, although highly connected interactors may outrank certain queries. Membrane-associated proteins are flagged and should be excluded from default dynamics simulations (i.e., free-in-solution) because they require separate modeling within a membrane environment and submission via the `traj` query (see Applications). Additional prioritization criteria could be incorporated at this stage (see Discussion). For example, pathways unique to bacterial or fungal species could be prioritized over those present in the human host; similarly, proteins from pathogenic organisms could be prioritized over those from commensal species, a distinction particularly important in the context of the human microbiome. Automating these criteria would require standard bioinformatics tools and workflows beyond the scope of this minimal implementation, but could improve the efficiency of systematic target identification.

In the second module (*cleaner*, *S_2_*), each predicted model from `dbac` or `sequ` is subjected to a sanitization process in which the full-length structure is partitioned into fragments by removing regions that fail to meet quality criteria. Contiguous sequences longer than 15 residues with per-residue confidence scores below 60 are removed, whereas shorter sequences are retained and treated tentatively as loops or linkers. Of the remaining fragments, only those containing 50 or more residues are advanced to the next stage, while shorter fragments are discarded and logged for review. Retained fragments are then evaluated for alignment errors. If two adjacent intra-fragment domains, separated by a retained low-confidence segment, have an inter-domain alignment error of ≤ 3 Å for at least three residue pairs, the segment is preserved as a genuine loop. Otherwise, the domains are considered independent, with the intervening segment functioning as a linker (see Applications). In such cases, the linker may be cleaved, and each domain treated as an individual fragment to improve simulation efficiency; however, this separation is not performed by default and is logged for user review. To unify terminology, proteins submitted via `coor` or `traj` are also referred to as fragments, although they are neither generated by nor subjected to the cleaning process and are treated as provided by the user.

Higher-order modules (*expanders*, *S_3_* – *S_M_*) process each fragment from *S_2_* to produce branch expansions based on pKa shifts or post-translational modifications (PTMs), generating multiple variants depending on their states, each treated independently as a new fragment. These modules are not yet automated, pending integration with state-specific databases (e.g., phosphoproteomics) or the availability of more reliable predictive methods (see *Discussion*).

Except for `traj` queries, which proceed directly to the analysis module, all fragments advance to the dynamics simulation stage. Each fragment is placed in a cubic box of pre-equilibrated water so that its center of mass (CoM) coincides with the box center. The fragment is rotated about its CoM so that its longest dimension (primary axis of inertia) aligns with one of the box’s major diagonals, minimizing box size; an image separation of ∼10 Å is imposed along with periodic boundary conditions in all three dimensions. Disulfide bonds are introduced for cysteine pairs whose sulfur atoms are separated by less than 3.75 Å. For `coor` and `traj` requests, protonation states are defined by the user, with different states requiring separate submissions; this is equivalent to the manual application of the *expanders* modules. For `sequ` and `dbac` queries, the default protocol assigns a charge of −1 to Asp and Glu and +1 to Arg and Lys, while histidine residues are treated as deprotonated. Sodium or chloride counterions are added at random in the solvent phase to neutralize the system’s net charge; by default, no additional salt is included. All proline residues are set to the *trans* configuration. Any transition to *cis* would require a different fragment, either generated by the *expanders* module or set manually and submitted through the `coor` or `traj` query. All simulation parameters can be modified via a master script, although the default settings are designed to simplify the workflow, enhance reproducibility, and reduce sensitivity to initial conditions.

Simulations are performed using NAMD with the CHARMM force field. Trajectories, including those provided by the user via `traj` requests, feed into the analysis module to compute metrics and associated data. Most quantities are calculated from statistically decorrelated snapshots, assumed here to be every 100 ps by default, based on typical decorrelation times for local properties. Analyses requiring higher temporal resolution, such as distance cross-correlations or the detection of transient water chains, use the full time series (every 2 ps) and thus take longer to compute. The default simulation length of 100 ns was chosen heuristically to ensure convergence of all metrics within a reasonable wall time (see documentation in [xxx] for instructions on modifying simulation parameters)).

### Metrics

The pipeline generates metrics derived from protein conformational dynamics, selected according to the following criteria: they must (i) quantify known or probable determinants of protein function; (ii) be computationally efficient; (iii) have a clear physical interpretation; (iv) be experimentally testable; (v) reflect known antimicrobial mechanisms of action (MoA); and (vi) serve as effective inputs for data-driven prediction (e.g., ML/AI models). Preference is given to local observables that converge rapidly over time. Based on a survey of known antimicrobial MoA (see Discussion), eight metrics are proposed, each defined by a three-letter code (Fig. 1C): surface topography (`TOP`); representative conformers and occupancies (`RCO`); local mobile elements (`LME`); local conformational fluctuations (`LCF`); polar (`PIN`) and nonpolar (`NIN`) interaction networks; distance-correlation networks (`DCN`); and internal water networks (`IWN`). These metrics can be supplemented with methods designed to identify potential catalytic sites.^29,30^ However, these additional metrics can be implemented as optional plugins rather than in the core workflow, since they do not require dynamics simulations and can, with varying levels of confidence, be predicted from a fixed structure (see Discussion).

In the definitions below, 〈… 〉_*t*_ denotes a time average. The index *k* refers to an atom belonging to residue *K*. *Ω*_*x*_(*t*) is the *local domain* of an atom *x*, defined as the set of atoms {*i*} of a specified type within a distance *d* of *x* at time *t*. *N*_*x*_(*t*) is the number of atoms *i* in this set. The temporal dependence of a quantity *Q* is generally omitted unless needed, with *Q*_0_ indicating its value at an initial time *t* = 0, after structural changes in the system have stabilized. Each metric is represented by a scalar quantity and stored in the `tempFactor` field (B-factor column) of the coordinate file of a reference conformation, enabling visualization as heat maps for comparison (see Applications and SI).

#### TOP

Surface topography is a quantity assigned to each atom *k*, used to distinguish concave and convex regions corresponding to surface crevices and protrusions. It is calculated as

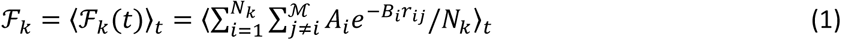

where *M* is the total number of atoms in the protein; *r*_*ij*_ ≡ |**r**_*i*_ − **r**_*j*_| is the distance between atoms *i* and *j* at time *t* (with **r** denoting atomic positions); and *A* and *B* are atom-dependent parameters.^31^ Thus, ℱ_*k*_ is a time-averaged measure of the degree of solvent exposure of atoms in the local domain of *k*. For surface atoms, ℱ captures the local concavity or convexity around *k*, expressed as a smoothed measure over the surface, allowing easy detection of crevices. Deep and narrow pockets often serve as binding sites for small molecules, whereas multiple adjacent pockets can accommodate specific chemical groups that may be linked to form larger, high-avidity ligands. These include peptides^32^ and polycyclic or branched molecules, which are common among antimicrobials. Some of these (e.g., rifamycins, macrolides, and streptogramins) target proteins, whereas larger compounds (e.g., enramycin, bacitracin, ramoplanin, and daptomycin) bind lipids or lipid-linked cell wall precursors. All are natural products, discovered either serendipitously or through systematic screening (see Introduction). Four related quantities are reported for each *k*: ℱ_*k*,*M*_ (maximum value of ℱ over the trajectory), ℱ_*k*,*m*_ (minimum), *σ*_*k*,ℱ_ (standard deviation), and Δ_*k*,ℱ_= ℱ_*k*,*M*_ − ℱ_*k*,*m*_ (range). Together, these quantities provide insight into the structure and dynamics of surface features, including the opening and closing of potential binding pockets and transient or hidden regions not apparent in static structures.

#### RCO

Conformational clustering is used to identify the most populated conformations adopted by a fragment during a simulation. Certain regions of the configurational space, defined here by the positions of all Cα atoms in the fragment, are sampled more frequently than others. Representative conformations corresponding to local density peaks in this space (clusters), referred to as conformers (*c*), and their occupancies (*p*), defined as the number of conformations within each cluster relative to the total number of snapshots considered from the trajectory, are identified using a density-based clustering algorithm. A *conformer* refers to a representative structure within a local density peak, around which a group of similar conformations is clustered. The conformation closest to the point of highest local density, defined as the number of neighboring points within a fixed radius *δ*, is selected first and designated as conformer *c*_1_. Similarity between conformations is assessed using the Cα-RMSD as a distance measure after rigid-body superposition of the Cα atoms that minimizes it. All conformations within a distance *δ* of *c*_1_are grouped into a conformational family centered on *c*_1_ and excluded from further analysis. The procedure is then repeated on the remaining unassigned conformations to identify the next densest region, represented by *c*_2_, and so on. This iterative process continues until all sampled conformations are assigned to one of *M* families, each associated with a distinct peak in the density landscape. The fragment’s dynamic behavior is thus described by a set of conformer–occupancy pairs {(*c*_*l*_, *p*_*l*_)}, where *l* = 1,2, … , *M* and ∑*p*_*l*_ = 1. By default, *δ* is set to 1.5 Å, and the first four and last four residues of the fragment are excluded from the analysis. All atoms of *c*_*l*_ are assigned the same *p*_*l*_ for visualization. Associated data include the `TOP` metric for each conformer, averaged over its local cluster.

#### LME

Local mobile elements identify local hinges, distortions, and changes in proximity, features that may have functional relevance (see Discussion). These elements are determined by

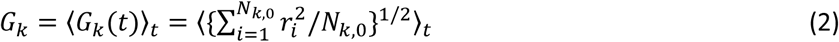

where *i* runs over all the atoms (*N*_*k*,0_) in the local domain of *k*. The domain is determined at an initial time *t*_0_, after artifactual conformational changes have subsided. Depending on the fragment, reaching this point can take several nanoseconds, but is heuristically set by default at 2 ns after full equilibration under NPT conditions. The C_α_ RMSD of the whole fragment can be monitored on the fly during the simulation using the command `rmsd` to ensure the choice of *t*_0_ is reasonable (see SI). The metric is instead intended to highlight local structural changes occurring after *t*_0_. All atoms of residue *I* are included if any atom of *I* is in Ω_*k*_(0). In Eq. 2, *r*_*i*_ = |*T***r**_*i*_ − **r**_*i*,0_|, where *T* is the roto-translational matrix that minimizes the RMSD between {**r**_*i*_} at time *t* and {**r**_*i*,0_} at *t*_0_; that is, the atoms in Ω_*k*_(0) are rigidly superimposed (oriented) onto the initial configuration before computing the summation over *i*. The metric thus ignores large scale movements. Only Cα atoms in Ω_*k*_ are considered, using *d* = 5 Å. For `LME` calculations, only C_α_ atoms are included in {*k*} and Ω_*k*_. A more general definition of *G*_*k*_ removes dependence on a single reference by moving *t*_0_ along the trajectory and averaging over *t*_0_, 〈*G*_*k*_〉_*t*0_. However, this approach is better suited to systems with large local conformational changes not expected on the time scales considered here. Any global conformational change^33^ must begin with local changes that are expected to be captured by this definition. Therefore, it is not necessary to characterize global motions to identify the local moiety as a targetable hotspot for drug binding. Larger-scale movements can be characterized using alternative metrics based on, for example, elastic network models,^34^ which may bypass the need for dynamical simulations.

#### LCF

Conformational fluctuations are defined by the standard deviations of *G*_*k*_(*t*) using suitable selection of atoms for summations and orientations,

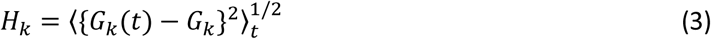

where *G*_*k*_(*t*) and *G*_*k*_ are defined in Eq. 2. Only side-chain atoms in Ω_*k*_ are used in the summation, while the Cα atoms in the domain define the orientation. The local environment of each residue is treated as a liquid patch subject to internal fluctuations, moving with the underlying structure and thus ignoring global motions. The metric is divided into side-chain atom fluctuations (`LCF`_sc_), where {*k*} and Ω_*k*_include all atoms except C_α_, and backbone fluctuations (`LCF`_bb_), where only C_α_ atoms are included. Ligands that bind proteins generally induce conformational rigidity, although in some cases they may increase flexibility. In both cases, these effects can alter protein function and, in pathogens, inhibit replication or growth (see Applications). Protein stability can be experimentally assessed in vitro and in living cells using thermal shift assays,^35–37^ which are increasingly used in early-stage drug development.

#### PIN and NIN

Polar and nonpolar interactions often form networks that may indicate structural vulnerability hotspots. Disruption of these networks, whether by mutations or ligand binding, can propagate through the protein and affect its function. Among polar interactions, hydrogen bonds and salt bridges are typically the strongest, although dipole–ion, dipole–dipole, and cation–π interactions can also contribute. `PIN` does not distinguish between these types but combines them into a single distance-based criterion to capture all relevant polar contacts; associated output data allow for deeper analysis. The number and frequency of contacts serve as a proxy for the interaction strength between a residue *K* and its neighboring residues {*K*_*i*_}. For the purpose of defining this distance criterion, all heavy atoms, excluding carbon, that are covalently bonded to a hydrogen atom are classified as donors (*D*), regardless of whether they participate in conventional hydrogen bonds. Acceptors (*A*) are defined as heavy atoms, excluding carbon, that can interact electrostatically with a donor. Residue *K* is considered to interact with residue *K*_*i*_ at time *t* if, for at least one *A* − *D* pair, the distance *δ* between *A* and the hydrogen atom bound to *D* is less than 3 Å. The output data also include a more stringent criterion, *δ* < 2 Å, to identify hydrogen bonds only. The total number of interactions for *K* at time *t* is 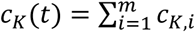 where *m* is the number of interacting residues and *c*_*K*,*i*_ is the number of interactions with *K*_*i*_. The overall interaction strength of residue *K* over the simulation is defined as *f*_*K*_ = ∫ *c*_*K*_(*t*)*dt*. This value is assigned to all atoms *k* ∈ *K*, so *f*_*k*_ = *f*_*K*_. The `PIN` metric is then *f*_*k*_ normalized by the maximum *f*_*K*_ in the fragment, measuring residue *K*’s local electrostatic contribution to structural stability relative to other residues. Raw, non-normalized values are also provided as associated data, facilitating downstream comparative analyses.

A similar criterion is used for the `NIN` metric. Nonpolar interactions can be conceptually grouped into dispersion attractions, π–π and CH–π interactions, aromatic–aliphatic contacts, and hydrophobic interactions in water-accessible regions such as the protein surface. These interaction types are not distinguished, but the output data allow for more detailed analysis. Instead, a carbon–carbon distance criterion (*δ*_*CC*_) is applied, with two side chains considered to engage in nonpolar interactions if *δ*_*CC*_ < 4.8 Å. The normalized metric *g*_*k*_ quantifies the overall strength of these interactions, whether they are purely nonpolar (e.g., buried interactions inaccessible to water and driven by dispersion forces) or hydrophobic (solvent-exposed contacts mediated by water). Raw values are also provided as associated data. Side chains of polar, acidic, or basic residues can also participate in nonpolar interactions through their aliphatic hydrocarbon groups, and the metric accounts for these contributions.

#### DCN

Distance correlations quantify both linear and nonlinear associations between random variables, in this case, atom-pair distances during a simulation.^38,39^ Unlike Pearson’s correlation, which measures linear relationships between individual atom pairs, distance correlations capture dependencies between entire motion patterns and equal zero only if the atoms move independently. This property enables the detection of complex, non-monotonic dependencies that Pearson’s correlation may miss. Distance correlations are calculated as

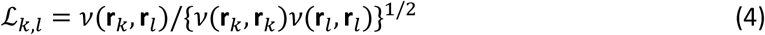

where *v*(**r**_*k*_, **r**_*l*_) is the distance covariance between atom positions **r**_*k*_ and **r**_*l*_, defined as *v*(**r**_*k*_, **r**_*l*_) = 〈〈*α*(*t*, *t*′)*β*(*t*, *t*′)〉〉_*t*,*t*′_, where *α*(*t*, *t*^′^) = *r*_*k*_(*t*, *t*^′^) − 2〈*r*_*k*_(*t*, *t*^′^)〉_*t*′_ + 〈〈*r*_*k*_(*t*, *t*^′^)〉〉_*t*,*t*′_ and *r*_*k*_(*t*, *t*^′^) ≡ |**r**_*k*_(*t*) − **r**_*k*_(*t*^′^)|. *β* is defined similarly by replacing *k* with *l*. The terms *v*(**r**_*k*_, **r**_*k*_) and *v*(**r**_*l*_, **r**_*l*_) follow from these definitions. The pipeline generates ℒ_*k*,*l*_ values for all (*k*, *l*) and also produces a subset representing long-range correlations only. By default, this subset includes the strongest correlations (ℒ > 0.9) between surface atoms, identified by their ℱ_*k*_ and ℱ_*l*_ (`TOP`) values, that are more than six positions apart in the primary sequence (approximately two helical turns) and spatially separated by at least one residue; spatial separation is enforced by requiring the C_α_ atoms of the two residues to be at least 7 Å apart. Additional output is generated using more stringent criteria (C_α_ atoms > 12 Å apart and ℒ > 0.99). These filters remove trivial correlations while retaining those likely relevant for long-range modulation (see Discussion). For each residue *K*, the maximum value of ℒ_*k*,*l*_ over all atoms *l* is identified and assigned to all atoms *k* ∈ *K* as the `DCN` metric. This value is then mapped onto the reference structure to enable visualization. By default, C_β_ atoms are used to characterize side-chain correlations (`DCN`_sc_) and C_α_ atoms for backbone correlations (`DCN`_bb_); the corresponding long-range metrics are also produced (`DCN`_sc,lr_ and `DCN`_bb,lr_). Raw ℒ_*k*,*l*_ values are reported as matrices and two-dimensional plots for further analysis. Pearson’s correlations and the corresponding plots are included in the output data for completeness.

#### IWN

Formation of water chains in the interior of proteins may have functional significance. Detecting hydrogen-bonded water chains requires high temporal resolution because they are typically short-lived, making this metric the most computationally demanding. The algorithm proceeds as follows. At each time point, all hydration water molecules are first extracted using a water–protein distance criterion. To identify buried water molecules from this subset, the degree of solvent exposure is calculated for each molecule using an analytical approximation.^31^ Molecules that are more than 70 % buried are considered internal to the fragment and retained, while the rest are discarded. The retained waters are then clustered^40,41^ using a definition in which two molecules are connected (i.e., hydrogen-bonded) if the distance between their oxygen atoms is less than 3 Å. For each trajectory frame, the number of dimers (*υ*_2_), trimers (*υ*_3_), …, *n*-mers (*υ*_*n*_) are recorded, along with the coordinates of the oxygen atoms in each *i*-mer and the coordinates of the fragment. All statistics in the metric-associated output are derived from these data. The metric `IWN` is a heat-map representation of the statistics of contacts between fragment residues and internal water clusters across the entire trajectory. This enables identification of regions that host internal water networks, including water chains capable of shuttling protons.^42^ The metric-associated data, including water-chain coordinates (`IWN`_o_) provide detailed information for further analysis.

## III. Applications

Some of these metrics have recently been applied to investigate molecular mechanisms beyond microbial systems, including human proteins,^42–45^ primarily through the use of the `traj` query following multiple independent simulations. The examples below illustrate the method’s potential for identifying targets in pathogens (see SI for additional command queries and the `examples/` directory in the distribution for other applications and further details).

### III.1. The method is first applied to a polyprotein of unknown structure using only its primary sequence. This example demonstrates the general analysis protocol for selecting potential hotspots with a mechanistic rationale

Human rhinovirus (HRV) cause upper respiratory infections, including common cold, and can exacerbate asthma and chronic pulmonary disease.^46^ They are classified into three species (A, B, and C), and no antiviral agents are currently approved for clinical use, although several compounds have been shown to inhibit viral replication in vitro, including vapendavir, the most recent candidate cleared as an investigational new drug.^47^

The HRV genome encodes a single polyprotein, a common feature of RNA viruses.^48^ This polyprotein is cleaved co- and post-translationally by the viral proteases 2A and 3C into four structural proteins (VP1–VP4), which form the capsid (60 copies each), and seven non-structural proteins involved in replication and host-cell modulation. Viral entry begins with receptor binding at the VP1 canyon, followed by endocytosis. Endosomal acidification triggers pH-dependent capsid conformational changes, uncoating, and RNA release into the cytoplasm for translation and replication.

Capsid variability largely determines the type of host receptor the virus uses for entry and its serotype specificity. Over 160 serotypes exist across the three species, reflecting antigenic diversity that has hindered the development of universal vaccines and broad-spectrum antivirals. The method is applied here to three HRV-B serotypes (strains 3, 14, and 72). The `<session>` file and complete output are provided in the SI. Because the AF database lacks viral sequences, polyprotein structures were generated using a standalone module (see Methods). Cleanup produced nine fragments, mostly corresponding to post-translational products (Fig. 2).

**Figure 2.**
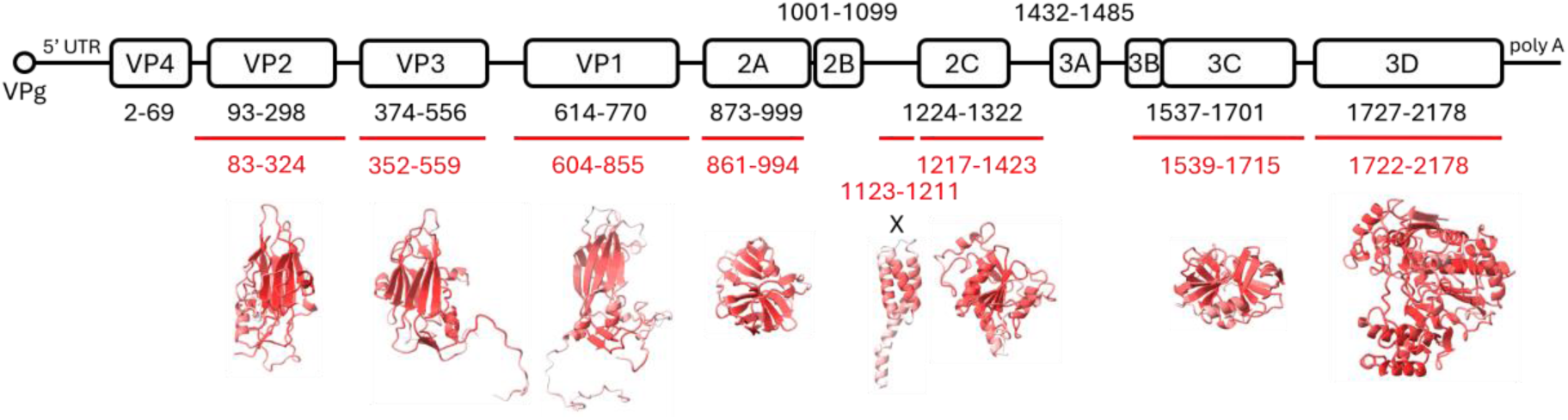
Human rhinovirus polyprotein encoded by the HRV-B3 genome, showing regions corresponding to structural proteins (VP1–VP4) and nonstructural proteins, including the membrane-associated protein 2B and the four catalytically active proteins 2A, 2C, 3C, and 3D. Predicted fragments (red) align closely with the expected proteins after polyprotein cleavage; however, the VP4 and 3A models were predicted with low confidence, and 2B was too short and was therefore excluded during preprocessing (see text). Fragment X was modeled with reasonable confidence and likely corresponds to the N-terminal region of 2C, despite not being explicitly annotated in the database entry (NCBI Protein: ABO69378.1).

VP1 is the best-characterized antiviral target; inhibitors typically stabilize one of the protein lobes, blocking uncoating and RNA release (Fig. S3A). 3C is a secondary target, while emerging targets include 3D and 2C. By contrast, VP2–VP4, 2A, 2B, 3A, and 3B are rarely pursued despite being potentially druggable, due to mechanistic constraints. For example, targeting the VP2/VP3 interface requires near-complete inhibition, whereas partial occupancy of VP1 inhibitors is sufficient to stabilize the capsid.

This example is not intended to propose new anti-HRV compounds, so no drug-development constraints are considered. Fragment X, the smallest, allows easier interpretation of most metrics, while VP2 and 3D provide complementary information.

The basic visualization output displays metrics across two screens (Fig. 3A). Associated text-based data are also included in the workflow output, allowing deeper analysis after visual inspection of regions of interest. The analysis both begins and ends with the `TOP` metric, as it highlights potential pockets and crevices, which are necessary features for ligand binding. The heatmap coloring (see Fig. 3 caption) makes the concavity and convexity of the molecular surface readily apparent. `TOP`_M_ and `TOP`_m_ highlight potential conformational changes during the dynamics, including short-lived or sporadic pocket opening and closing events. The magnitude of these changes can be visualized through `TO`P_σ_ and further examined quantitatively through explicit analysis of pocket conformations and occupancies once a pocket of interest is identified (see Analysis below). The extent of the variable regions should serve as a minimal criterion for defining a binding hotspot in any downstream analysis, whether for high-throughput screening or de novo design.

**Figure 3.**
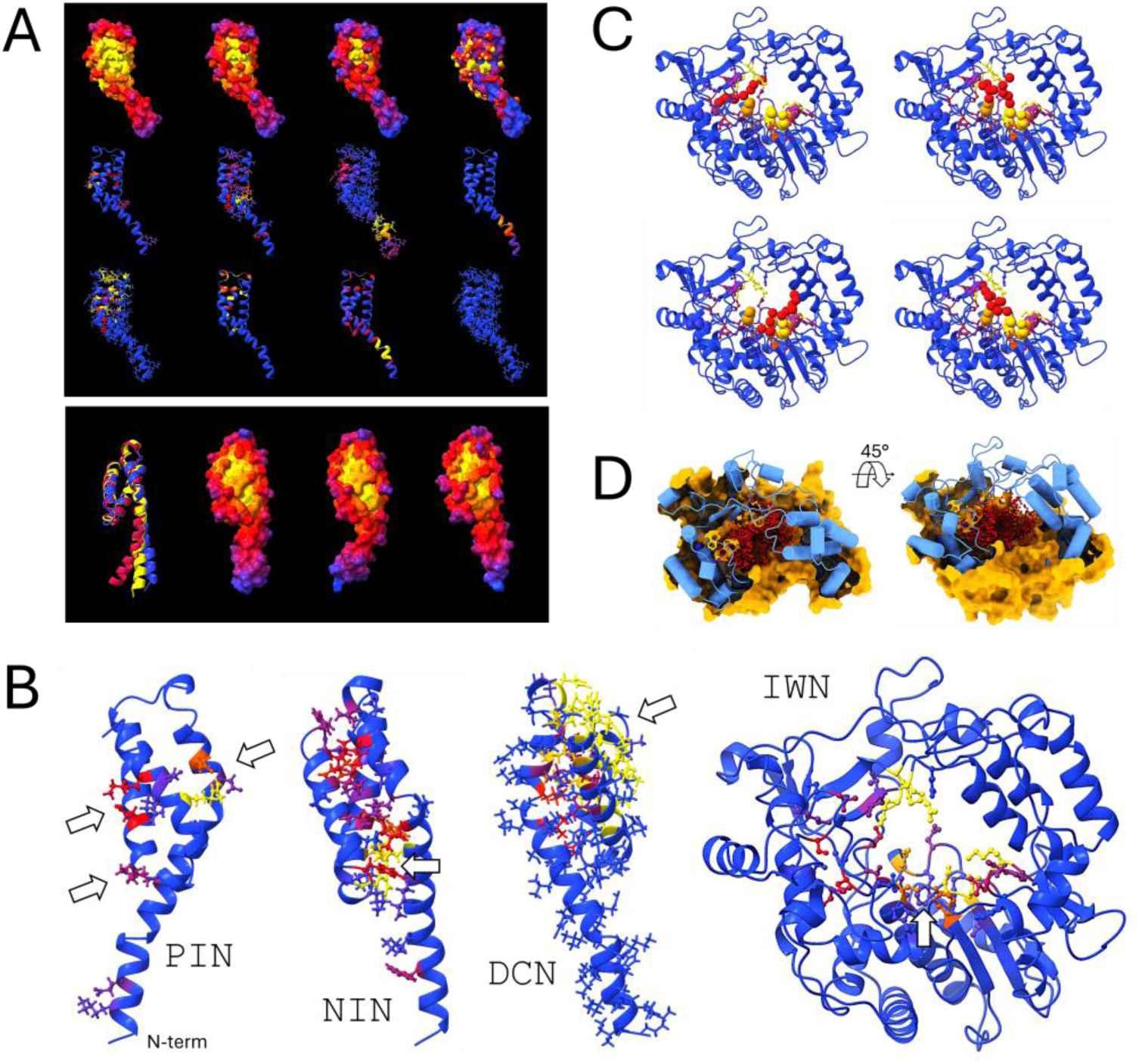
Metrics visualization. (A) Default metric distributions for each fragment (shown for X of HRV serotype B3; see Fig. S1 for VP2 and 3D). Upper screen, left to right: `TOP`_M_, `TOP`, `TOP`_m_, `TO`P_σ_ (upper row); `PIN`, `NIN`, `LCF`_sc_, `LCF`_bb_ (middle); `DCN`_sc_, `DCN`_bb_, `LME`, `IWN` (lower). Lower screen, left: `RCO` (> 5 %, Cα-superimposed; three conformers shown colored by occupancy); right: `TOP` of main conformers. When present, `IWN` produces an additional screen showing water-chain configurations (not observed in fragment X), and `DCN`-associated plots and matrices are also generated as output data (Fig. S1 for VP2 and 3D). (B) Selected metric details highlight features of interest (see text), with `IWN` shown for fragment 3D, the only fragment with buried water chains. Colors are chosen to enhance heatmap contrast (blue, low; red, intermediate; yellow, high). Panels rotate independently about the fragment center to facilitate metric comparison and hotspot identification. `TOP`, `PIN`, `NIN`, and `IWN` are normalized per fragment; absolute values are provided in the output data for comparative analysis. Typical heatmap ranges are 20–80 (blue <20, yellow >80). `LCF` and `LME` are reported in Å and are comparable across fragments, with recommended ranges of 0.5–1.0 and 1.0–2.0, respectively. Only the highest `DCN` values are mapped, with heatmap ranges from 0.95 to 0.99. `RCO` conformers are colored by occupancy (10–80%). All ranges are indicative, heuristically determined, and used as defaults. (C) Buried water networks are labeled `IWN`_o_ (index refers to water oxygen; non-empty IWM metric only for fragment 3D, as shown). Four water decamers for serotype B14 are shown as red spheres, highlighting cationic and anionic residue contacts in the central cavity. (D) Aggregation of all water decamers observed in the simulation; fragment shown with backbone in blue and part of the surface removed. Four aspartates relevant for metal-ion coordination during catalysis are shown as van der Waals spheres (hydrogens omitted). Water clusters range from dimers to >50-mers, with size distributions, populations, and coordinates provided in the output data. Analysis of fragment 3D reveals multiple crevices lining both the internal cavity and outer surface, suggesting potential allosteric effects of ligand binding on catalysis and protein–protein interfaces, respectively (see SI).

Fragment X shows four crevices at the interfaces of adjacent helices within the crown-like three-helix bundle (Fig. 2). In the absence of additional information, all four pockets are legitimate targets a priori, presenting deep or extended topographic depressions. Additional metrics are useful for assessing the potential effects of ligand binding in each pocket. `PIN` highlights three regions where internal hydrogen bonding or ionic interactions stabilize the protein surface (Fig. 3B); disruption of these local, solvent-accessible clusters could have important structural consequences. `NIN` suggests that nonpolar interactions stabilize the core of the crown (Fig. 3B), with two phenylalanine residues at the base playing a central role. However, these nonpolar clusters are internal and thus unlikely to be affected by a surface-targeting ligand. Statistics of `PIN` and `NIN` pairwise interactions are provided as plain-text files for further analysis, and scripts for generating plots are included in the distribution.

`LCF`_sc_ and `LCF`_bb_ indicate that the protein is largely rigid, except near the tip of the N-terminal helix, which is unlikely to represent a valuable target. This lack of flexibility suggests that the opening and closing of crevices indicated by `TOP`_M_ and `TOP`_m_ are likely transient and sporadic. Quantitatively, the standard deviation of these fluctuations is less than 0.6 Å. Regions of high local rigidity or mobility indicate features worth exploring as potential hotspots for ligand binding. `DCN`_sc_ (Fig. 3B) shows that correlations are mostly restricted to the tip of the crown. However, `DCN`_bb_ analysis, supported by the long-range `DCN`_bb,lr_ plot (not shown; see Fig. S1 for VP2 and 3D), indicates correlated movements across different sides of the crown. This suggests that ligand binding on one side could influence the other. `LME` shows that local backbone distortions are also restricted to the N-terminal helix, while `RCO` (Fig. 3A, lower panel) indicates that these distortions primarily involve local backbone hinges triggered by a proline. The `IWN` metric shows that there are no buried water chains. This metric is important because it may reflect chemical rather than structural functions, and it is observed only in fragment 3D, the RNA polymerase. It also appears in fragments studied in sections III.2 and III.3, in both cases associated with known catalytic activity. For fragment 3D, analysis of `IWN` (Fig. 3B) reveals numerous polar and ionic side chains lining the internal cavity and in contact with hydrogen-bonded water clusters; the heatmap coloring reflects contact statistics. These residues include three aspartates involved in metal ion coordination, as well as others that contribute to substrate positioning, mediate proton transfer, and stabilize reaction intermediates. In this case, water molecules do not act as nucleophiles in the polymerization reaction but may play different roles before, during, and after catalysis. Associated output data also include statistics on water cluster sizes and populations, indicating that these clusters range from dimers to over 50-mers. Analysis of `IWN`_o_ (Fig. 3C, showing decamers) demonstrates that the water networks crisscross the cavity, often connecting cationic residues in the upper lobe with the catalytic basin, and are spatially concentrated on the protein’s inner surface (Fig. 3D). This metric underscores the importance of carefully considering desolvation effects when evaluating ligand binding, particularly when using simplified scoring functions.

The analysis above indicates that the four crevices identified through TOP are located in regions capable of influencing various metrics to differing degrees. Three of these pockets may disrupt two or more interaction networks, and one additionally affects long-range residue cross-correlations. Therefore, ligand binding in these regions may not only interfere with the fragment’s interactions with other proteins or the membrane but also produce intrinsic structural and functional effects. Identifying such regions and their potential impacts is the primary goal of the metric analysis.

The method is designed to scan the proteome, or subsets of it, without prior knowledge of the functional roles of protein fragments. For the system considered here, information on mutations across the three HRV-B serotypes can be used to focus on the pockets most likely to provide broad-spectrum antimicrobial targets. Automation of this prioritization could be implemented in the *sorter* module. Fragment X is ∼88 % identical across the three serotypes. Based on mutation positions, two of the four potential binding sites identified above are fully conserved and are mapped onto serotype B3 in Figs. 4 and S2 (residues in green, left panel). `TOP` analysis shows that one of these concave regions (Fig. 4) is larger with relatively little internal restructuring during the dynamics, except at the periphery of the crevice (`TO`P_σ_); the other is a smaller pocket (Fig. S2) showing transient and sporadic opening–closing dynamics, as suggested by the `LCF` analysis above. Binding to the larger crevice does not directly affect any of the metrics; aside from potential allosteric effects, its primary impact would likely be on interactions with other proteins or, for this fragment, with lipid molecules. Although binding to the smaller pocket could have a similar effect on interactions with other molecules, it may also disrupt one of the local hydrogen-bond networks or alter long-range side-chain correlations.

**Figure 4.**
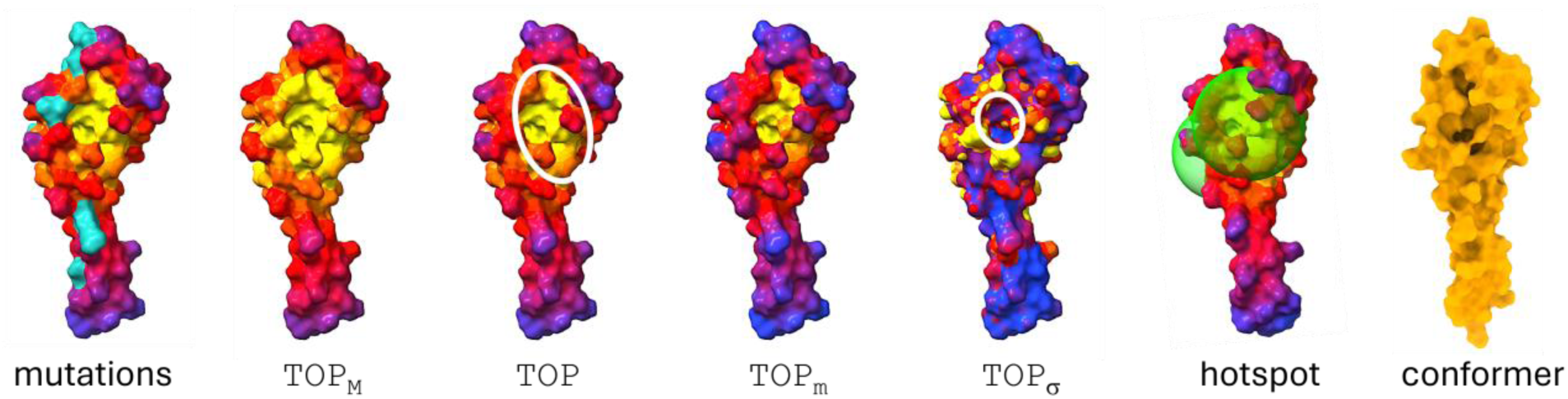
One of four potential ligand-binding sites in fragment X of HRV serotype B3. From left to right: mutations across the three serotypes (cyan); expanded yellow regions in `TOP`_M_ indicate pocket shrinkage (reduced solvent exposure or increased concavity) during dynamics; average topography (`TOP`) reveals the extent of the region suitable for ligand binding; color changes in `TOP`_m_ (reduced yellow, increased red–blue) indicate pocket opening (increased solvent exposure or convexity); local topography variability (`TO`P_σ_) shows minimal changes in the pocket interior (white circle) and greater changes at the periphery. These features help define the residues that need to be considered for hotspot clustering: a spherical domain (green) is used to cluster pocket conformations, where the second sphere corresponds to a secondary pocket (see text and Fig. S2). The molecular surface of the single hotspot conformer at full occupancy indicates that `TOP` variations arise primarily from side-chain movements, highlighting the need to account for side-chain flexibility in downstream analyses and ligand design.

Once the sites of interest are selected, their internal structures can be assessed. The implementation allows searching for local conformers and occupancies of each site using a clustering method similar to that employed in RCO via the command:

`$ ./get_hotspot_RCO.sh <hotspots>`

Here,

`<hotspots>` (e.g., `targets.hsp`) is an unformatted plain-text file containing any number of lines, each representing a hotspot defined by its center (`hspc`) and its size (`hspr`), with an optional comment field:

`hspc hspr ! (comments)`

where `hspc` is the residue number of the Cα atom considered closest to the pocket center, and `hspr` is the radius (in Å) of a sphere encompassing the pocket, the extent of which can be inferred from the `TOP` analysis as discussed above. These values are not set automatically and must be determined manually after visual inspection of the metric. For the larger and smaller pockets considered here, the regions are shown as green spheres in Fig. 4, and the corresponding `targets.hsp` file is provided in the SI. Unlike the `RCO` metric, which applies to fragments, the clustering radius for identifying hotspot conformers is set to 1 Å, as conformers separated by a larger Cα RMSD are treated as independent hotspots in downstream analyses. Although separating hotspots into multiple conformers allows the backbone to remain fixed in molecular docking methods, side-chain flexibility is still assumed to be necessary. This requirement can be cautiously avoided if the clustering explicitly includes side chains. In that case, the clustering threshold should be considerably lower, or false negatives are almost certain. For each pocket in fragment X, only a single conformer with nearly 100 % occupancy was observed (Figs. 4 and S2, rightmost panel), indicating that conformational changes associated with variations in local topography arise mainly from side-chain movements.

An example of multiple hotspot conformers is provided by a similar analysis of VP2 (Fig. S3). Metric analysis identifies five major crevices located at interfaces with VP3, VP1, and VP4 in the mature capsid (Fig. S3A), while four additional hotspots are non-interfacial but may still influence these interfaces allosterically (see Discussion and SI). As with fragment X, suitable ligands binding to these pockets can affect several metrics, in this case `PIN`, `NIN`, `LCF`, `LME`, and `DCN` (Fig. S1). It is reasonable to expect that the more properties a ligand targets, the greater its potential deleterious effect on protein function. The relevance of each metric can be explored systematically by assessing the impact of point mutations through dynamic simulations or by repeated application of the method to the mutant followed by comparative metric analysis.

To identify hotspots relevant to ligands affecting all three serotypes, site conservation was assessed. For fragment VP2, serotypes B3 and B14 are identical, whereas B72 shares only 87 % sequence identity with either. Four conserved pockets were thus identified: one with potential effects on VP2 or VP4 assembly (Fig. 5), and three at interfaces with VP1, VP2, and VP3 in the mature capsid (Fig. S3B). `TOP` analysis indicates that these four pockets have internal structural variability. Hotspot clustering (see `targets.hsp` in SI) shows between one and three conformers, depending on the hotspot. For the hotspot in Fig. 5, conformer c_1_ might be expected to be the most effective target, but c_2_, despite its lower population in the free fragment, may offer higher affinity or avidity for a suitable ligand. Both conformers must therefore be considered, with side-chain flexibility taken into account for each. The pipeline provides a number of scripts for downstream analysis, particularly for de novo design of cyclic peptides. This was used to target c_1_ in VP2 as an illustration, and the results are available in the SI and in the `examples/` directory of the distribution.

**Figure 5.**
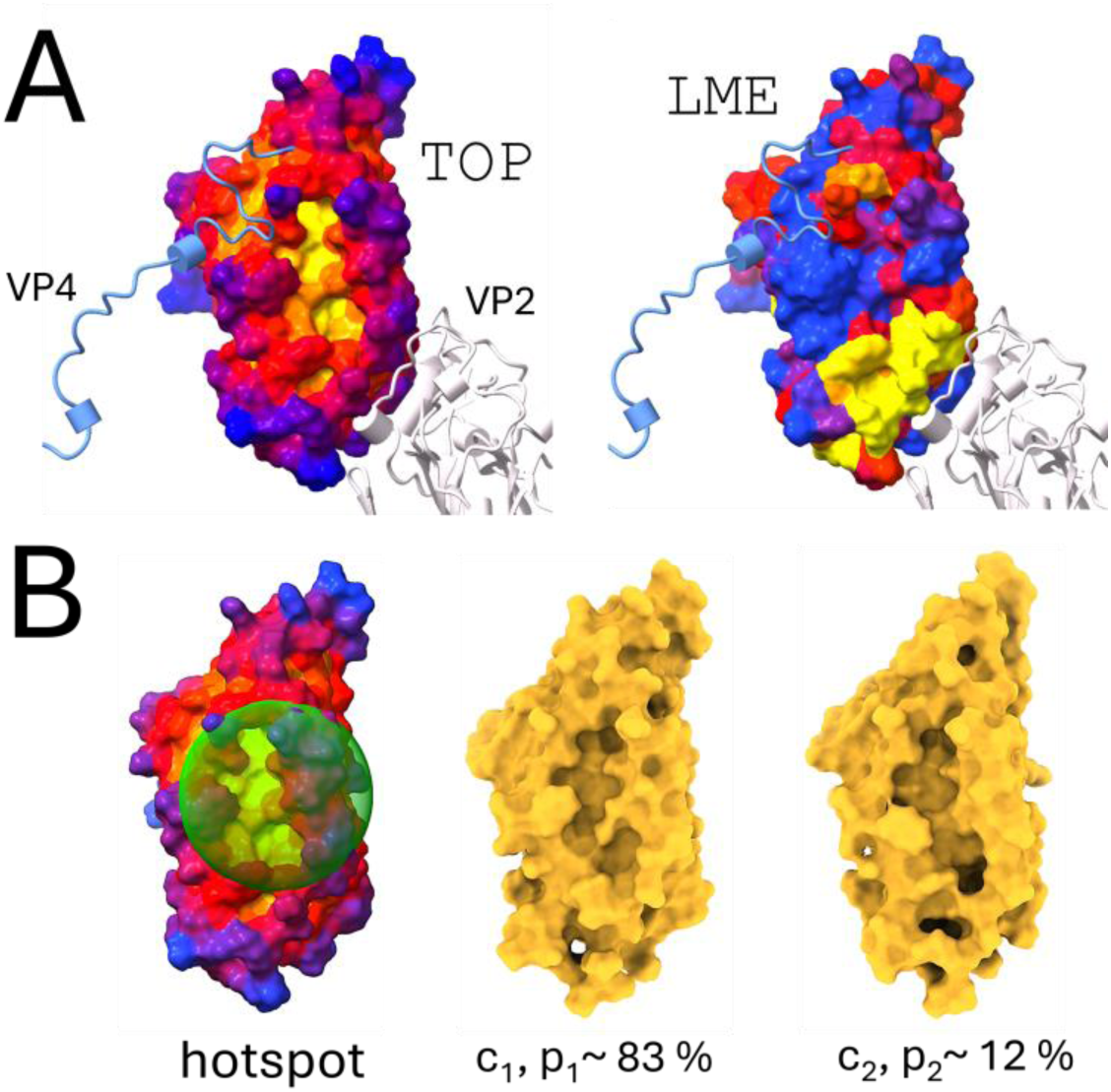
A) Potential allosteric pocket on fragment VP2 of HRV serotype B14 (left), affecting VP4 binding and neighboring VP2 assembly in the growing capsid. A mobile loop (yellow, right) separates the pocket from the VP2–VP2 interface (see Fig. S1). (B) A spherical region is used to define pocket conformers and occupancies, revealing two states in the free fragment determined by local backbone conformations (see text). This pocket, along with three others at protein–protein interfaces that presumably affect capsid assembly (see Fig. S3), is conserved across the three serotypes. Each conformer must be treated as a separate fragment, and side-chain flexibility must be explicitly included in all downstream analyses. An example of de novo design of cyclic peptides targeting both conformers is provided in the SI to illustrate the use of a class of output scripts.

### III.2. The method was applied to metabolic networks to evaluate its ability to detect known drug targets and identify potential new ones

The mycobacterial inner membrane contains transporters that shuttle molecules from the cytosol to the periplasm, an intermediate step in cell-wall assembly. The exporter MmpL3 translocates trehalose monomycolate (TMM) in *M. tuberculosis* (Mtb), contributing to outer mycomembrane assembly. Because of its essential role, MmpL3 is a promising anti-tuberculosis target, as inhibiting TMM efflux disrupts cell-wall formation and compromises bacterial viability. Some metrics have recently been used to propose a mechanism of efflux.^42^ However, MmpL3 is only one of many proteins involved in wall biosynthesis, lying at the intersection of multiple pathways in mycolic acid–arabinogalactan–peptidoglycan (mAGP) biosynthesis (Fig. S4). The mAGP complex^49^ forms a three-component scaffold that constitutes the cell-wall envelope, producing the thick, waxy, and impermeable barrier responsible for resistance to antibiotics and host defenses. These pathways, like MmpL3, sustain cell-wall synthesis and provide alternative therapeutic targets that can be systematically explored with the proposed method.

Two metabolic pathways are considered: arabinogalactan biosynthesis and mycolic acid biosynthesis^50,51^ (Fig. S4). The former involves cytosolic and inner-membrane–associated enzymes, while the latter includes membrane, periplasmic, and secreted enzymes. Depending on localization, enzymes may be targeted by small compounds (cytosolic) or polycyclic/branched peptides and mimetics (extracellular). Modeling membrane-associated enzymes requires the membrane environment and a separate `traj`-query submission, which is not done here (see section III.3 for a demonstration in another system).

Figure 6 presents the networks generated by the query with interactors included (see session submission in Supporting Information). Although all queried proteins and their interactors are potential targets, the analysis focuses on highly connected nodes and cytosolic or periplasmic proteins. Three proteins of interest in pathway mtu00572 and six in mtu00074 are highlighted. Two proteins in mtu00572 (UniProt AC: P9WNL7 and P9WNL9) are membrane-associated enzymes and established targets of ethambutol, which inhibits arabinogalactan transfer into the mycobacterial cell wall. The third protein (P9WQF3) is a cytosolic carrier identified as a potential drug target despite limited characterization. Because it interacts with the other two proteins and the original request (see SI) did not include dynamics simulations for interactors, no metrics were generated for it; it is therefore not considered further but could be resubmitted as a standalone `unip` query for metric analysis.

**Figure 6.**
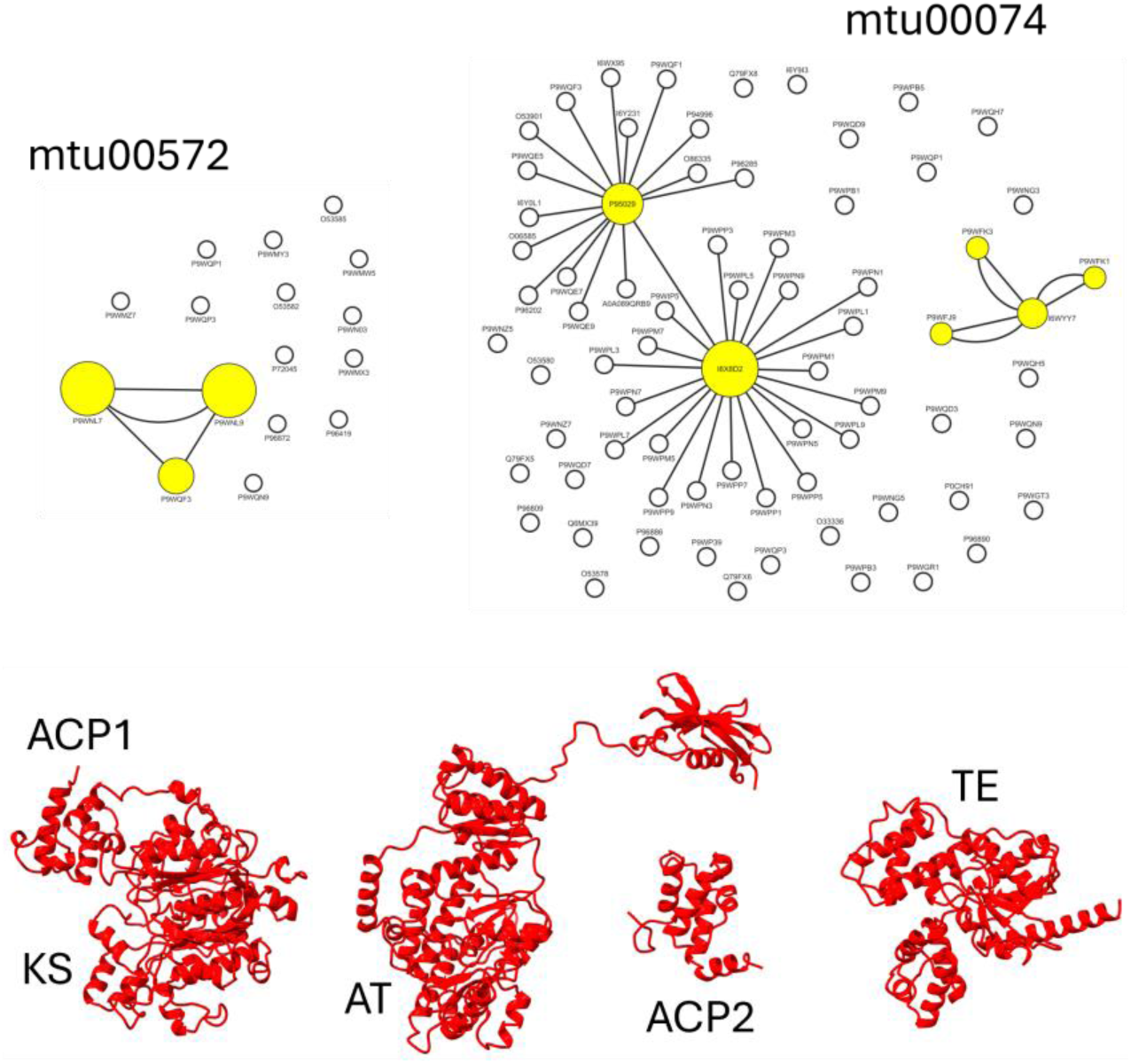
From network to binding sites. Upper panel: Two networks involved in mycolic acid–arabinogalactan–peptidoglycan (mAGP) biosynthesis (Fig. S4): mycolic acid biosynthesis (KEGG ID: mtu00074) and arabinogalactan biosynthesis (mtu00572). UniProt accession numbers are indicated; see Fig. 1 for details. The query retrieved interactors, i.e., proteins experimentally shown to be part of the same physical complexes as the queried proteins at some point during the cell cycle. Highly connected proteins are colored yellow; their perturbation may hinder mAGP synthesis, and most correspond to known or suspected antimicrobial targets. The protein analyzed here, Pks13 (I6X8D2), is a cytoplasmic enzyme for which only a functional domain (TE) has been validated as a viable antituberculosis drug target (see Fig. 7). Lower panel: Fragments of Pks13 (UniProt accession: I6X8D2): ACP1 (N-terminal acyl carrier protein), KS (ketoacyl synthase), AT (acyltransferase), ACP2 (second acyl carrier protein), and TE (thioesterase). The four fragments identified by the method encompass these functional domains; cryo-EM structures are available for some (8CV1, 8CUY) but were not used in this application (see the corresponding query in the SI).

In mtu00074, two proteins (P9WFJ9 and P9WFK1) are localized to the plasma membrane and are not known drug targets. A large multidomain protein (P95029) contains both membrane-associated and cytosolic domains and has been tentatively linked to pretomanid and pyrazinamide. Another protein (P9WFK3) localizes to the plasma membrane and peptidoglycan cell wall and is not a known drug target. Protein I6WYY7 has no defined localization or known drug associations, whereas I6X8D2 is a cytoplasmic protein with a domain inhibited by coumestan derivatives and is analyzed further in this example.

Polyketide synthase Pks13^52,53^ is a 1,733-residue protein comprising five main domains, with the thioesterase (TE)-like C-terminal domain, corresponding to fragment 4 in Fig. 6 (lower panel), serving as the primary target of known inhibitors.^54^ Each fragment presents features that could be perturbed to affect function, but only fragment 1 (ACP1-KS) is analyzed here. The results are presented succinctly, as the analysis follows the general protocol described in III.1. Fragment 1 has three main conformers with approximate occupancies of 68 %, 18 %, and 7 % (see the visualization package provided as Suppl. Data). Analysis of the metrics (Fig. S5A) suggests three potential hotspots. One corresponds to a deep crevice (Fig. S5B, middle panel) that harbors the catalytic site, including residues C287, H423, and H463 (numbering refers to the full-length protein). This region is highly rigid, as indicated by `LCF` and `LMB`. Metrics `PIN` and `IWN` further suggest that a ligand could exploit or disrupt local clusters of hydrogen-bonded residues and water chains, which span assemblies from dimers to 30-mers. As revealed by `IWN`_O_, many of these networks are located near the catalytic residues and crisscross the crevice (Fig. S6).

A second, smaller pocket is observed at the ACP1/KS interface (Fig. S5B, left). Assuming ACP1 and KS are not cleaved in or around the DE-rich linker (residues ∼76–92), ligand binding in this region is predicted to affect both `PIN` and `NIN`, the latter including F79, a residue for which the F79S mutation has been reported to confer resistance.

Finally, multiple shallow pockets are present on the outer surface of the KS globular domain (Fig. S5B, right), which is also relatively rigid. `DCN` analysis indicates that this region contains numerous cross-correlated residues and may also act as an interface with other proteins.

Once a potential target is identified, the method can be applied in a more focused manner through coor or traj queries, using an experimentally determined structure or trajectories from more detailed modeling and simulations performed separately (see III.3). Identifying a highly promising target may also motivate experimental structure determination and biochemical and biophysical studies of its function and potential strategic value as a viable druggable target.

### III.3. The method was applied to multiple independent simulations of a monomeric protein to illustrate how converged metrics can be obtained. The example also examines a protein complex to show how nontrivial hotspots, emerging only in the context of the complex’s dynamics, can be identified within each monomer

*Staphylococcus aureus* D-alanine:D-alanine ligase (Ddl) is a cytosolic enzyme that catalyzes the ATP-dependent ligation of two D-alanine molecules to form D-ala–D-ala, a key precursor in peptidoglycan biosynthesis.^55,56^ Peptidoglycan, a complex polymer forming the structural backbone of the bacterial cell wall, provides rigidity and shape; it is thick and exposed at the cell surface in Gram-positive bacteria, whereas in Gram-negative bacteria it is thin and situated within the periplasmic space. Catalysis requires ATP and D-ala as substrates, with Mg^2+^ as an essential cofactor and K^+^ enhancing enzymatic activity. Owing to its central role in cell-wall formation, Ddl is an attractive but underexploited antimicrobial target. Although no clinically approved drugs directly inhibit this enzyme, several preclinical inhibitors have been reported, including allosteric compounds that induce conformational changes and suppress catalytic activity.^57^ Additional inhibitor classes have demonstrated *in vitro* activity but remain limited in potency.^54,58,59^

Available crystallographic data for both apo and ligand-bound forms (PDB IDs: 2I87 and 7U9K) reveal a homodimeric arrangement. Complexes with co-crystallized allosteric inhibitors (PDB ID: 2I80) highlight regulatory sites that induce conformational shifts and may serve as alternative drug-binding sites. This dimeric architecture appears conserved across bacterial species, although it is unclear whether dimerization is essential under native conditions. This question is addressed in this example as a byproduct of comparative metric analysis. If dimerization is required for activity, disrupting dimer formation could compromise complex stability and function, representing a potential antimicrobial strategy. Studying both monomeric and dimeric forms can therefore guide the design of compounds that destabilize the dimer, prevent its assembly, or directly affect catalysis. Both forms were considered, with three independent simulations performed for the monomer to illustrate convergence.

Because several loops are missing in all available experimental structures, the monomer was obtained from the AF2 database using the `unip` query (see the corresponding request in the SI), and the dimer was generated using an in-house AF2-Multimer implementation. Structural alignment with the highest-resolution crystal structure (PDB ID: 2I87) showed that the experimentally resolved regions of both models have a Cα RMSD of ∼1.5 Å, which is within the expected range of structural fluctuations under physiological conditions and within the thresholds used for the `RCO` calculations.

Multiple simulations can be necessary for unknown systems, such as fragments emerging from querying networks (see III.2), where fragment behavior may depend strongly on initial conditions, which are not known a priori. In this application, three independent simulations were performed for the monomer. Each simulation was intentionally brief to highlight incomplete sampling and the resulting variability in the computed metrics; combining an increasing number of simulations demonstrates convergence. Robust statistics are obtained only after convergence within and between simulations, so three to five simulations of 100 ns each may be sufficient in most cases, since all metrics are local properties. Metrics were first calculated using simulation replicas 1 and 2, and then recalculated after adding replica 3, showing progressive convergence. Simulations are combined using the following command:

`$ ./combine.sh <requests>`

Here,

`<requests>` specifies which replicas to combine (e.g., `monomer_12.cmb` or `monomer_123.cmb`, for combination of 1 and 2, or 1, 2, and 3, respectively). Each

`<requests>` script consists of a single plain-text line in the form:

[request_id]_1_ [request_id]_2_ … [request_id]_N_ > [combout] ! (comments)

where `[request_id]`_i_ denotes the name of request `i` in the `<session>` files(s), and `[combout]` specifies the name for the combined results. The corresponding commands for the monomer are provided in the SI. The output is written to a dedicated directory, `combined.requests.[combout]/`. The term “combination” is used rather than “average” because not all metrics are computed as means over simulation replicas (ensemble); for example, `TOP`_M_ and `TOP`_m_ are defined as the maximum and minimum values over the aggregated ensemble (that is, across replicas), and `DCN` is treated as a normalized average. The metrics show differing sensitivities to sampling: `TOP` is the least sensitive, whereas `DCN` (Fig. S7) and `LCF` (Fig. 7A,B) are generally the most sensitive, although convergence can be achieved with only a few replicas.

**Figure 7.**
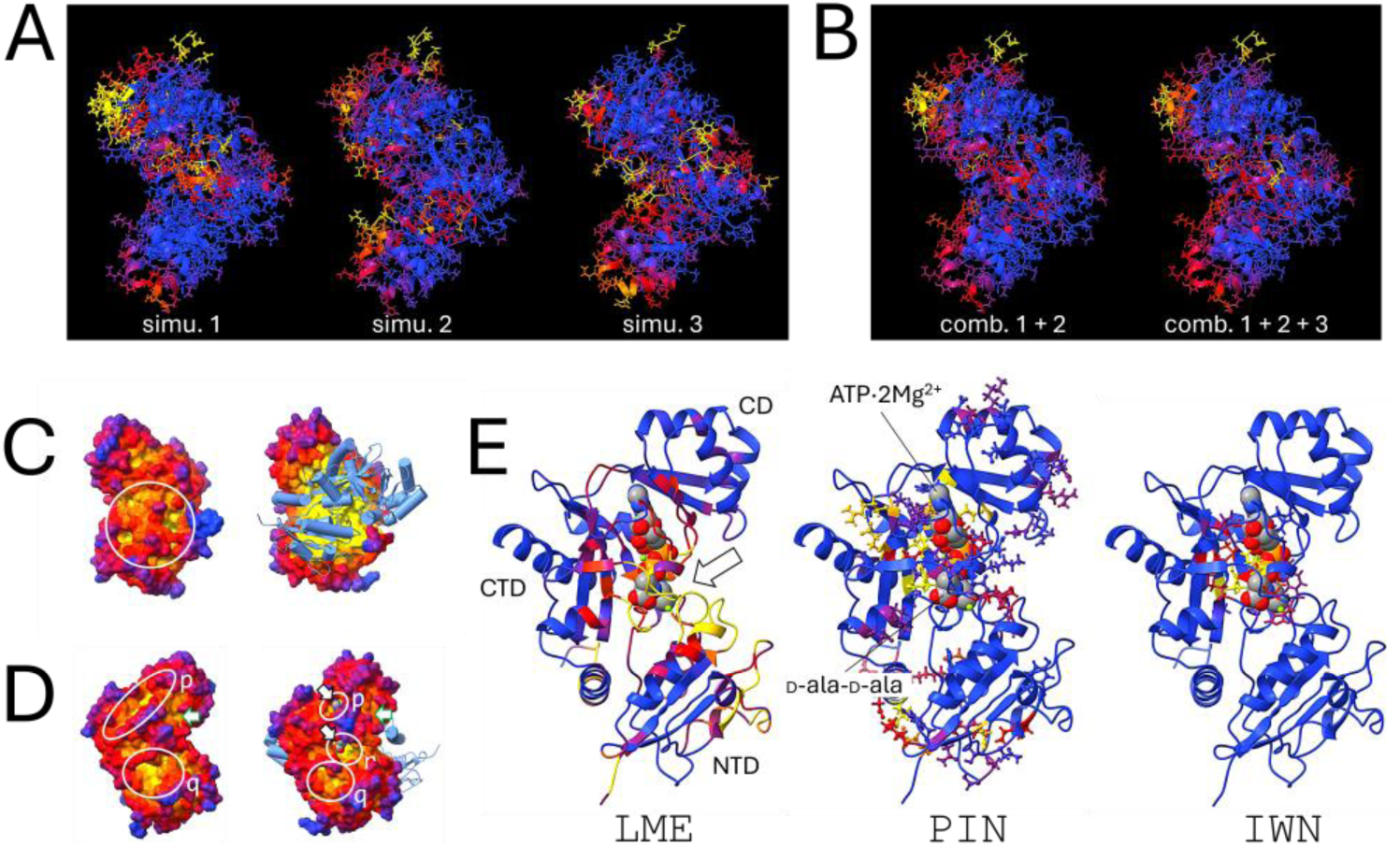
Metric analysis for the monomer and dimer of *S. aureus* D-alanine:D-alanine ligase (Ddl). (A) Metric dependence on simulations and convergence: `LCF` from three independent simulations of the monomer; simulations were intentionally short to highlight differences. (B) Gradual convergence is achieved by combining an increasing number of simulation replicas. Some metrics are more sensitive to initial conditions and trajectories than others; `DCN` provides another example of convergence through replica combination (see Fig. S7). (C) `TOP` view of the free (left) and homodimerized monomer (right; backbone of the second monomer shown in light blue), highlighting potential binding hotspots at the dimerization interface (white circle; see also Fig. S8). (D) Opposite face of the structures shown in (C), illustrating potential hotspots (circled) that appear and disappear (r) or change shape (p and q) depending on whether the monomer is free or dimerized. An alternate pocket, partially occluded in the view, leads to the same internal cavity as p and is indicated by green arrows (pocket s in Fig. S8C). The arrow pointing to p indicates the ATP molecule, and the arrow pointing to r indicates the D-Ala-D-Ala substrate from an *E. coli* Ddl dimer co-crystal, which has a similar overall fold and was backbone-superimposed (both ligands are rendered as van der Waals spheres). (E) Three features identified by the converged metric analysis of the free monomer, anticipated to be modulated by ligand binding to the hotspots, all located within the region containing the catalytic cleft. NTD, N-terminal domain; CD, central domain; CTD, C-terminal domain.

Metric analysis identifies multiple potential hotspots in the uncomplexed Ddl monomer. Some of these are further examined in the context of the dimer to assess how dimerization affects monomer behavior and their potential as drug targets both in the unbound state and within the complex. This analysis addresses an aspect not explored in Applications III.1 or III.2.

Without prior knowledge of the monomer’s biological function, the metrics identify several hotspots that, a posteriori, are recognized as functionally critical. Figure 7C presents the `TOP` metric for the free (left column) and dimerized (right) monomer. Several small pockets and crevices are apparent in the circled region, corresponding to the dimerization interface. These pockets expand and contract during the dynamics, as indicated by `TOP`_M_ and `TOP`_m_ (Fig. S8A), although these changes are sporadic and short-lived, reflecting the relative rigidity inferred from `LME` and `LCF` (Fig. 7B,E). Nevertheless, transient or rare conformations may still be exploited for ligand binding (cryptic sites) and are more effectively captured by a focused pocket conformer analysis (see III.2).

Analysis of the remaining metrics indicates that ligand binding to this region would not disrupt `PIN` or `NIN`, while `DCN` reveals communication largely confined to the smaller lobe (Fig. S7; N-terminal domain, NTD, in Fig. 7E), where no significant exploitable pockets are detected. Collectively, these observations suggest that ligands targeting this region in the uncomplexed monomer would primarily exert structural effects by perturbing dimer formation. Dimerization may also be perturbed by targeting the interface boundary of a preformed dimer, as revealed by `TOP` analysis (Fig. S8B), where transient crevices could serve as wedging sites. A striking example of this mechanism is ivermectin, which binds at the interface of the glutamate-gated chloride channel and stabilizes it in an open state, ultimately leading to parasite death.^60^

Analysis of the opposite side of the free monomer (Fig. 7D) reveals markedly different behavior. Several pockets and crevices differ both structurally and dynamically between the free monomer and its dimeric form. Notably, crevice p in the monomer becomes a deeper but smaller pocket upon dimerization; the large pocket q present in the monomer is reduced in the dimer; and, most significantly, a deep pocket r emerges in the dimer, being only weakly apparent in the free monomer. These rearrangements arise from hinging of the three domains (NTD, CD, and CTD; Fig. 7E) relative to one another, as well as from reshaping of loops at their interfaces (arrow in Fig. 7E) upon dimerization. The resulting local structural changes, comprising two loops in the NTD, one in the CD, and two in the CTD, all in close proximity, are highlighted by the `LME` metric (Fig. 7E). Analysis of co-crystal structures from other species (e.g., 5C1P, 4C5C)^61,62^ shows that pockets p and r correspond to the binding site for ATP, D-ala–D-ala, and the required Mg^2+^ ions and water molecules involved in substrate orientation and catalysis (Fig. 7D, right).

Although the `TOP` metric alone identifies all these regions as potential binding sites, metrics `LME`, `PIN`, and `IWN` indicate that ligand binding can induce local conformational changes, perturb extended hydrogen-bond networks, or disrupt water-mediated pathways in or around the catalytic site. Consequently, unlike ligands targeting the dimerization interface, binding to these pockets is likely to produces functional rather than structural effects. Binding to pocket q, in particular, may alter `LME` in a manner that allosterically affects substrate binding. Targeting this pocket in both the monomer and dimer, potentially with distinct ligands, could maximize disruption. This could be achieved by stabilizing the closed conformation of pocket r either before or after dimerization, in both cases impairing substrate access. Alternatively, direct targeting of pockets p and r would orthosterically inhibit catalysis by interfering with ATP or substrate binding. An alternate pocket leading to the same internal cavity as p lies to the right of the panels (indicated by the green arrows; see Fig. S8C). In all cases, `LME` suggests allosteric perturbation, `PIN` indicates disruption of hydrogen-bond networks stabilizing the CTD and CTD/CD interface, and `IWN` points to altered water and ion accessibility essential for coordination during catalysis, consistent with crystallographic observations of ordered water molecules and ions in complexes with ATP or ADP. Binding to these pockets is likely synergistic and dependent on the binding order, which may complicate lead design or identification when multiple pockets are targeted simultaneously. In this case, each protein–ligand complex should be queried independently, for example using the `traj` query, and metric analysis should be performed for further characterization.

## IV. Discussion

Pathogens evade antimicrobials through various mechanisms, most commonly via point mutations that alter drug targets. Genetic adaptations, such as horizontal gene transfer or gene duplication and divergence, provide alternative escape routes. Well known mechanisms include^63–69^ enzymatic drug inactivation (e.g., β-lactamases), reduced permeability (loss or modification of porins), and active efflux via drug export pumps. Some strategies redirect essential pathways: bypassing a drug target, where the original enzyme remains functional but an alternative enzyme or pathway compensates; or inactivating a drug-sensitive enzyme and replacing its function through another mechanism. Additional defenses include target protection, where proteins shield the drug target without altering it; biofilm formation, which limits drug penetration and promotes tolerance; and persistence, where dormant subpopulations survive treatment and can repopulate later.

Moreover, microorganisms possess an inherent adaptability to gene inactivation, making the lasting efficacy of a single compound, or even a few in combination, unlikely.^70,71^ In *S. cerevisiae*, more than 80% of single-gene knockouts do not severely impair growth,^72^ yet many combinations of two or three non-essential gene deletions lead to synthetic lethality, showing that organismal robustness can be overcome.^73,74^ Similar observations occur in other pathogens, including *M. tuberculosis* and related mycobacteria, where perturbing multiple non-essential genes can lead to bacterial death, even though single knockouts alone do not.^75–77^

Given the broad repertoire of microbial self-preservation mechanisms, it is unrealistic to expect that one or a few compounds could produce lasting clinical effects. The strategy proposed here shifts the focus from identifying lead compounds, a task already well supported by computational methods, to the less-explored but highly desirable goal of identifying multiple targets with therapeutic potential. This framework emphasizes protein dynamics rather than static structures. While the methods used in this integration have limitations, including force fields, databases, and knowledge-based predictors, the strategy relies on the reasonable expectation that ongoing advances will provide more reliable and efficient tools. Its design ensures continued relevance amid rapid technological and methodological developments and is unlikely to require frequent adaptations.

Biological networks are generally robust to random disruptions but sensitive to targeted perturbations of central nodes. Experiments and simulations show that removing nodes with high degree or betweenness centrality compromises network integrity,^24–28^ identifying these proteins as key vulnerabilities that, in pathogens, can impair survival or growth. Combining network-vulnerability analysis^78^ with target-site characterization allows strategic prioritization and can guide antimicrobial discovery. Building on this principle, the workflow scans targets across multiple biological levels, from genus and species to networks, pathways, genes, proteins, and their physical interactors, extending to statistically significant conformers of hotspot regions. The concept of polypharmacology,^19–21^ originally defined as the action of a single drug on multiple targets, has been broadened to include the coordinated action of multiple agents, effectively generalizing the traditional notion of combination therapy. The method and workflow proposed here provide a systematic computational approach to address this challenge.

The core idea of the method is a set of metrics capturing structural and dynamic features important for protein function and frequently targeted by antimicrobials. For example, adamantane derivatives block proton transfer in influenza A virus by inhibiting H^+^ transport through M2 channels, a key step in viral uncoating, forming the basis of amantadine and rimantadine.^79^ A similar mechanism has been proposed for Mtb, where proton transfer through the MmpL3 transporter drives the movement of mycolic-acid precursors across the inner membrane.^42,80^ Bedaquiline, a diarylquinoline, also inhibits Mtb ATP synthase by disrupting proton transport.^81,82^ Hydrogen-bond networks in neuraminidase have been linked to influenza pandemic strains^83^ and other antimicrobial activity.^84,85^ Increased flexibility in viral spike proteins, such as SARS-CoV-2, enhances infectivity^86–88^ while decreased flexibility of capsid proteins blocks rhinovirus infectivity (see III.1). Local structural features, including hinge regions, may further support replication efficiency, as reported for respiratory syncytial virus^89^ and SARS-CoV-2.^90^ Similar conformational elements are targeted by antimicrobials: salinamide disrupts the helix–loop transition of the RNA-polymerase trigger loop in *E. coli*;^91,92^ ivermectin wedges open glutamate-gated chloride channels,^60,93^ causing unregulated ion flow in parasites; and fidaxomicin hinders hinge movement in the Mtb RNA-polymerase clamp.^94^ Regardless of the mechanism, molecular topography must provide suitable binding sites, either deep crevices for small, high-affinity ligands or clusters of pockets for larger, high-avidity compounds.

Therefore, all the metrics, despite their simplicity, are directly related to relevant mechanisms of action. Distance correlation may have more limited practical relevance, but it is retained because it provides a cohesive view of dynamics and may help identify potential allosteric sites or mechanisms, which is a key goal of the method. While many antimicrobials target enzyme active sites, allosteric inhibition is increasingly recognized as a selective alternative. Throughout this paper, the term is used broadly to refer to the modulation of any protein function through long-range conformational changes, not limited to effects on catalytic sites or enzyme activity. Clinically relevant examples targeting membrane-bound proteins include echinocandins, which bind a noncatalytic site in fungal β-(1,3)-D-glucan synthase, blocking glucan polymerization;^95,96^ bedaquiline, which allosterically modulates mycobacterial ATP synthase;^97,98^ and maraviroc, which prevents HIV entry by binding the CCR5 receptor.^99^ Soluble or cytosolic examples include fidaxomicin, which inhibits bacterial RNA polymerase near the RNA exit channel,^100,101^ and rifampicin, which stabilizes a conformation blocking RNA elongation;^102^ an alternative mechanism has also been proposed.^103^ Other drugs, such as pyrazinamide and certain InhA inhibitors, modulate enzyme conformation to reduce activity.^104,105^ Notably, no antifungal agents currently act through allosteric mechanisms, although structural studies suggest that soluble fungal enzymes, such as aldolases, may harbor potential allosteric sites. To date, such mechanisms remain largely unexplored. Fungal pathogens are emerging as a major threat to public health, including *Pneumocystis*, *Candida*, and *Aspergillus* species. Only four classes of antifungal drugs are currently available, underscoring the urgent need for new therapeutic strategies.^106–108^ A promising application of the method is antifungal target discovery against resistant strains.

Allosteric sites are difficult to identify a priori, but the workflow is designed to facilitate their detection. Exploiting allosteric modulation provides additional treatment options, in which drugs can act either directly or as adjuncts to enhance the effects of other agents. Allosteric inhibitors may also reduce the likelihood of resistance compared with active-site inhibitors. Single mutations can abolish binding of orthosteric inhibitors without impairing enzyme function, whereas allosteric inhibitors target regulatory sites that control conformation and dynamics. As a result, overcoming allosteric inhibition generally requires multiple coordinated mutations that counteract ligand-induced conformational and dynamic changes while preserving catalytic activity, making resistance less likely.

The workflow’s preprocessing module should account for changes in protonation states, cis–trans isomerization of peptide bonds, particularly at X–Pro motifs, and post-translational modifications (PTMs), all of which play central roles in microorganisms. Each modified and unmodified form should be treated as a separate fragment generated by the expander modules. For example, pH varies across organs, tissues, and cellular compartments and can change over time, typically ranging from 5 to 8, although it may be lower in specific microenvironments; these variations can influence microbial growth and antibiotic efficacy.^109–111^ Accurate pKa prediction remains challenging, and although many computational approaches have been proposed, their integration is deferred to future work due to current limitations.

Similar considerations apply to PTMs.^112^ Protein function can be altered either by modifying the PTM site or by disrupting interactions with the enzymes that catalyze it.^113^ PTM disruption may destabilize cellular functions and has been implicated in diverse pathologies.^114^ Disrupting these processes can compromise tightly regulated cellular functions, making PTM-targeting strategies potentially effective against pathogens. Hundreds of PTM types have been identified in microorganisms, affecting nearly all canonical amino acids.^115^ While some are rare, transient, or species-specific (e.g., bacterial two-component phosphorylation systems), the most prevalent include phosphorylation, acetylation, methylation, ubiquitination, glycosylation, and proteolytic cleavage.^116^ Except for proteolysis, which is irreversible, these PTMs are dynamically reversible. Therefore, both unmodified and modified fragments should be included in the workflow and treated on equal footing. Their integration is also deferred to future work.

Finally, oncology provides a notable example of the types of applications targeted by the proposed method, where features such as allostery, cryptic-site identification, and polypharmacology have been central to recent advances in the development of KRAS inhibitors and their emerging relevance to pancreatic cancer.^117^ Cancer cells and microbial pathogens share the capacity for adaptive evolution under therapeutic pressure. Although cancer arises from host cells whereas microbial pathogens are independent organisms, both systems have analogous evolutionary dynamics that drive resistance. The development of KRAS inhibitors illustrates how a historically ‘undruggable’ protein was rendered tractable through the discovery of an exploitable allosteric pocket, particularly in KRAS G12C-mutant tumors.^118,119^ First-generation inhibitors such as sotorasib and adagrasib established the clinical feasibility of this approach but were largely limited to specific mutations and frequently undermined by rapid resistance evolution. More recently, second-generation therapeutics such as daraxonrasib,^120^ a multi-selective RAS inhibitor, have expanded this strategy toward a broader spectrum of KRAS-driven cancers, raising the possibility of more durable pathway suppression.

## Conclusions

The proposed method provides a simplified and standardized framework for antimicrobial target discovery as a precursor to lead identification. It is grounded in the premise that coordinated perturbation of biological networks using multiple compounds at submaximal doses may reduce toxicity and limit mutational escape. At its core, the approach relies on a reduced set of dynamics-based metrics that capture key features of known antimicrobial mechanisms. The framework is extensible across biological scales, from single proteins to whole proteomes, enabling systematic multi-target strategies for antimicrobial discovery. It is implemented as a lean, modular workflow optimized for portability, reproducibility, and minimal user intervention, and built entirely on standard open-source components and Linux toolchains for deployment in both institutional high-performance computing environments and local workstations.

## Supporting information

The supplemental file includes a first page serving as an index describing its contents

## Acknowledgments

All simulations and analyses were conducted on the NIH Biowulf HPC cluster (https://hpc.nih.gov/), with additional validation on an HPC cluster at the Texas Advanced Computing Center (TACC; https://tacc.utexas.edu/); individual modules were tested in a high-throughput computing (HTC) environment via the Open Science Grid (OSG; https://osg-htc.org/). A prototype of this pipeline, including documentation, a tutorial, and examples, has been deposited in the Open Science Framework (DOI: 10.17605/OSF.IO/EM2WH). The author thanks the Office of Cyber Infrastructure and Computational Biology, NIAID, NIH.

## References

1 Berdy, B., Spoering, A. L., Ling, L. L. & Epstein, S. S. In situ cultivation of previously uncultivable microorganisms using the ichip. Nat. Protoc. 12, 2232–2242 (2017).

2 Ling, L. L. et al. A new antibiotic kills pathogens without detectable resistance. Nature 517, 455–459 (2015).

3 Murray, C. J. L. et al. Global burden of bacterial antimicrobial resistance in 2019: a systematic analysis. The Lancet 399, 629–655 (2022).

4 Robbins, N., Caplan, T. & Cowen, L. E. Molecular Evolution of Antifungal Drug Resistance. Annual Review of Microbiology 71, 753–775 (2017).

5 Mehta, K. et al. Modernizing Preclinical Drug Development: The Role of New Approach Methodologies. ACS Pharmacology & Translational Science 8, 1513–1525 (2025).

6 Kadri, S. S. et al. Difficult-to-Treat Resistance in Gram-negative Bacteremia at 173 US Hospitals: Retrospective Cohort Analysis of Prevalence, Predictors, and Outcome of Resistance to All First-line Agents. Clinical Infectious Diseases 67, 1803–1814 (2018).

7 Taubenberger, J. K., Powers, J. H. & Bhattacharya, J. The new vision from the National Institute of Allergy and Infectious Diseases (NIAID). Nature Medicine 32, 791–792 (2026).

8 Warr, W. A., Nicklaus, M. C., Nicolaou, C. A. & Rarey, M. Exploration of Ultralarge Compound Collections for Drug Discovery. Journal of Chemical Information and Modeling 62, 2021–2034 (2022).

9 Neumann, A., Marrison, L. & Klein, R. Relevance of the Trillion-Sized Chemical Space “eXplore” as a Source for Drug Discovery. ACS Medicinal Chemistry Letters 14, 466–472 (2023).

10 Mullowney, M. W. et al. Artificial intelligence for natural product drug discovery. Nature Reviews Drug Discovery 22, 895–916 (2023).

11 Melo, M. C. R., Maasch, J. R. M. A. & de la Fuente-Nunez, C. Accelerating antibiotic discovery through artificial intelligence. Communications Biology 4, 1050 (2021).

12 Fernandes, P. Antibacterial discovery and development—the failure of success? Nature Biotechnology 24, 1497–1503 (2006).

13 Brötz-Oesterhelt, H. & Sass, P. Postgenomic Strategies in Antibacterial Drug Discovery. Future Microbiology 5, 1553–1579 (2010).

14 Brown, E. D. & Wright, G. D. Antibacterial drug discovery in the resistance era. Nature 529, 336–343 (2016).

15 Payne, D. J., Miller, L. F., Findlay, D., Anderson, J. & Marks, L. Time for a change: addressing R&D and commercialization challenges for antibacterials. Philosophical Transactions of the Royal Society B: Biological Sciences 370, 20140086 (2015).

16 McDowell, L. L., Quinn, C. L., Leeds, J. A., Silverman, J. A. & Silver, L. L. Perspective on Antibacterial Lead Identification Challenges and the Role of Hypothesis-Driven Strategies. SLAS Discovery 24, 440–456 (2019).

17 Butler, M. S. et al. A Review of Antibacterial Candidates with New Modes of Action. ACS Infectious Diseases 10, 3440–3474 (2024).

18 Corre, C. et al. Discovery of Late Intermediates in Methylenomycin Biosynthesis Active against Drug-Resistant Gram-Positive Bacterial Pathogens. J. Amer. Chem Soc. 147, 40554–40561 (2025).

19 Haldar, J. Combination Therapy: A Pillar in the Fight Against Infectious Diseases. ACS Infectious Diseases 11, 3379–3385 (2025).

20 Cichońska, A., Ravikumar, B. & Rahman, R. AI for targeted polypharmacology: The next frontier in drug discovery. Current Opinion in Structural Biology 84, 102771 (2024).

21 Kabir, A. & Muth, A. Polypharmacology: The science of multi-targeting molecules. Pharmacological Research 176, 106055 (2022).

22 Zasloff, M. Antimicrobial peptides of multicellular organisms. Nature 415, 389–395 (2002).

23 Hassan, S. A. & Steinbach, P. J. Modulation of free energy landscapes as a strategy for the design of antimicrobial peptides. Journal of Biological Physics 48, 151–166 (2022).

24 Iyer, S., Killingback, T., Sundaram, B. & Wang, Z. Attack Robustness and Centrality of Complex Networks. PLOS ONE 8, e59613 (2013).

25 Zhang, X. et al. In silico Methods for Identification of Potential Therapeutic Targets. Interdisciplinary Sciences: Computational Life Sciences 14, 285–310 (2022).

26 Albert, R., Jeong, H. & Barabási, A.-L. Error and attack tolerance of complex networks. Nature 406, 378–382 (2000).

27 Barabási, A.-L. & Oltvai, Z. N. Network biology: understanding the cell’s functional organization. Nature Reviews Genetics 5, 101–113 (2004).

28 Alon, U. An Introduction to Systems Biology: Design Principles of Biological Circuits. Second edn, (Chapman and Hall/CRC, 2019).

29 Petrova, N. V. & Wu, C. H. Prediction of catalytic residues using Support Vector Machine with selected protein sequence and structural properties. BMC Bioinformatics 7, 312 (2006).

30 Song, J. et al. PREvaIL, an integrative approach for inferring catalytic residues using sequence, structural, and network features in a machine-learning framework. Journal of Theoretical Biology 443, 125–137 (2018).

31 Hassan, S. A. & Steinbach, P. J. Water-exclusion and liquid-structure forces in implicit solvation. J. Phys. Chem. B 115, 14668 (2011).

32 Chatterjee, S. Special Issue: Peptide Therapeutics. Journal of Medicinal Chemistry 69, 6297–6297 (2026).

33 Grimaldo, M., Roosen-Runge, F., Zhang, F., Schreiber, F. & Seydel, T. Dynamics of proteins in solution. Quarterly Reviews of Biophysics 52, e7 (2019).

34 Zhang, H., Gur, M. & Bahar, I. Global hinge sites of proteins as target sites for drug binding. Proceedings of the National Academy of Sciences 121, e2414333121 (2024).

35 Celej, M. S., Montich, G. G. & Fidelio, G. D. Protein stability induced by ligand binding correlates with changes in protein flexibility. Protein Science 12, 1496–1506 (2003).

36 Llowarch, P., Usselmann, L., Ivanov, D. & Holdgate, G. A. Thermal unfolding methods in drug discovery. Biophysics Reviews 4, 021305 (2023).

37 Almqvist, H. et al. CETSA screening identifies known and novel thymidylate synthase inhibitors and slow intracellular activation of 5-fluorouracil. Nature Communications 7, 11040 (2016).

38 Székely, G. J., Rizzo, M. L. & Bakirov, N. K. Measuring and testing dependence by correlation of distances. Ann. Statist. 35, 2769–2794 (2007).

39 Roy, A. & Post, C. B. Detection of Long-Range Concerted Motions in Protein by a Distance Covariance. Journal of Chemical Theory and Computation 8, 3009–3014 (2012).

40 Hassan, S. A. Computational Study of the Forces Driving Aggregation of Ultrasmall Nanoparticles in Biological Fluids. ACS Nano 11, 4145 (2017).

41 Hassan, S. A. Microscopic Mechanism of Nanocrystal Formation from Solution by Cluster Aggregation and Coalescence. J. Chem. Phys. 134, 114508 (2011).

42 Lee, Y.-S., Hassan, S. A. & Hurt, D. E. Proton Transfer Mechanisms in the MmpL3 Transporter of Mycobacterium tuberculosis Studied by Computer Simulations. ACS Bio & Med Chem Au DOI: 10.1021/acsbiomedchemau.5c00274 (2026).

43 Collette, A. M. et al. An unusual dual sugar-binding lectin domain controls the substrate specificity of a mucin-type O-glycosyltransferase. Science Advances 10, eadj8829 (2024).

44 Kumari, P. et al. Structural mechanism of CB1R binding to peripheral and biased inverse agonists. Nature Communications 15, 10694 (2024).

45 Iyer, M. R. et al. Unlocking Selenium Chemical Space via a Programmable Synthesis Platform Bearing Cannabinoid Receptor Recognition Motifs. J. Amer. Chem Soc. 148, 18649 (2026).

46 Jacobs, S. E., Lamson Daryl, M., St. George, K. & Walsh Thomas, J. Human Rhinoviruses. Clinical Microbiology Reviews 26, 135–162 (2013).

47 Lanko, K. et al. Comparative analysis of the molecular mechanism of resistance to vapendavir across a panel of picornavirus species. Antiviral Research 195, 105177 (2021).

48 Yost, S. A. & Marcotrigiano, J. Viral precursor polyproteins: keys of regulation from replication to maturation. Current Opinion in Virology 3, 137–142 (2013).

49 Alderwick, L. J., Harrison, J. K., Lloyd, G. S. & Birch, H. L. The Mycobacterial Cell Wall—Peptidoglycan and Arabinogalactan. Cold Spring Harb. Perspect. Med. 5, a021113 (2015).

50 Jankute, M., Grover, S., Rana, A. K. & Besra, G. S. Arabinogalactan and Lipoarabinomannan Biosynthesis: Structure, Biogenesis and Their Potential as Drug Targets. Future Microbiology 7, 129–147 (2012).

51 Jankute, M., Cox, J. A. G., Harrison, J. & Besra, G. S. Assembly of the Mycobacterial Cell Wall. Annual Review of Microbiology 69, 405–423 (2015).

52 Kim, S. K. et al. Structure and dynamics of the essential endogenous mycobacterial polyketide synthase Pks13. Nature Structural & Molecular Biology 30, 296–308 (2023).

53 Krieger, I. V. et al. SuFEx-based antitubercular compound irreversibly inhibits Pks13. Nature 645, 755–763 (2025).

54 Xia, F. et al. Targeting polyketide synthase 13 for the treatment of tuberculosis. European Journal of Medicinal Chemistry 259, 115702 (2023).

55 Suzuki, Y. et al. D-alanine synthesis and exogenous alanine affect the antimicrobial susceptibility of Staphylococcus aureus. Antimicrobial Agents and Chemotherapy 69, e01936–01924 (2025).

56 Qin, Y., Xu, L., Teng, Y., Wang, Y. & Ma, P. Discovery of novel antibacterial agents: Recent developments in D-alanyl-D-alanine ligase inhibitors. ChemMedChem 98, 305–322 (2021).

57 Liu, S. et al. Allosteric inhibition of Staphylococcus aureus d-alanine:d-alanine ligase revealed by crystallographic studies. Proceedings of the National Academy of Sciences 103, 15178–15183 (2006).

58 Becker, R., Pederick, J. L., Dawes, E. G., Bruning, J. B. & Abell, A. D. Structure-guided design and synthesis of ATP-competitive N-acyl-substituted sulfamide d-alanine-d-alanine ligase inhibitors. Bioorganic & Medicinal Chemistry 96, 117509 (2023).

59 Triola, G. et al. ATP competitive inhibitors of d-alanine–d-alanine ligase based on protein kinase inhibitor scaffolds. Bioorganic & Medicinal Chemistry 17, 1079–1087 (2009).

60 Huang, X., Chen, H. & Shaffer, P. L. Crystal Structures of Human GlyRα3 Bound to Ivermectin. Structure 25, 945–950.e942 (2017).

61 Tran, H.-T. et al. Structure of d-alanine-d-alanine ligase from Yersinia pestis: nucleotide phosphate recognition by the serine loop. Acta Crystallographica Section D 72, 12–21 (2016).

62 Batson, S. et al. Inhibition of D-Ala:D-Ala ligase through a phosphorylated form of the antibiotic D-cycloserine. Nature Communications 8, 1939 (2017).

63 Darby, E. M. et al. Molecular mechanisms of antibiotic resistance revisited. Nature Reviews Microbiology 21, 280–295 (2023).

64 Elbaiomy, R. G., et al. Antibiotic Resistance: A Genetic and Physiological Perspective. MedComm 6, e70447 (2025).

65 Belay, W. Y. et al. Mechanism of antibacterial resistance, strategies and next-generation antimicrobials to contain antimicrobial resistance: a review. Frontiers in Pharmacology Volume 15 - 2024 (2024).

66 Urban-Chmiel, R. et al. Antibiotic Resistance in Bacteria—A Review. Antibiotics 11, 1079 (2022).

67 Ahmad, M., Aduru, S. V., Smith, R. P., Zhao, Z. & Lopatkin, A. J. The role of bacterial metabolism in antimicrobial resistance. Nature Reviews Microbiology 23, 439–454 (2025).

68 Smith, W. P. J., Wucher, B. R., Nadell, C. D. & Foster, K. R. Bacterial defences: mechanisms, evolution and antimicrobial resistance. Nature Reviews Microbiology 21, 519–534 (2023).

69 Shepherd, M. J. et al. Ecological and evolutionary mechanisms driving within-patient emergence of antimicrobial resistance. Nature Reviews Microbiology 22, 650–665 (2024).

70 Kim, S., Lieberman, T. D. & Kishony, R. Alternating antibiotic treatments constrain evolutionary paths to multidrug resistance. Proceedings of the National Academy of Sciences 111, 14494–14499 (2014).

71 Schmid, A. et al. Monotherapy versus combination therapy for multidrug-resistant Gram-negative infections: Systematic Review and Meta-Analysis. Scientific Reports 9, 15290 (2019).

72 Giaever, G. et al. Functional profiling of the Saccharomyces cerevisiae genome. Nature 418, 387–391 (2002).

73 Costanzo, M. et al. A global genetic interaction network maps a wiring diagram of cellular function. Science 353, aaf1420 (2016).

74 Rahiminejad, S., De Sanctis, B., Pevzner, P. & Mushegian, A. Synthetic lethality and the minimal genome size problem. mSphere 9, e00139–00124 (2024).

75 Kalia, N. P. et al. Exploiting the synthetic lethality between terminal respiratory oxidases to kill Mycobacterium tuberculosis and clear host infection. Proceedings of the National Academy of Sciences 114, 7426–7431 (2017).

76 Lun, S. et al. Synthetic Lethality Reveals Mechanisms of Mycobacterium tuberculosis Resistance to β-Lactams. mBio 5, 10.1128/mbio.01767–01714 (2014).

77 Xu, Y., Ehrt, S., Schnappinger, D. & Beites, T. Synthetic lethality of Mycobacterium tuberculosis NADH dehydrogenases is due to impaired NADH oxidation. mBio 14, e01045–01023 (2023).

78 Holme, P., Kim, B. J., Yoon, C. N. & Han, S. K. Attack vulnerability of complex networks. Physical Review E 65, 056109 (2002).

79 Takizawa, N. & Yamasaki, M. Current landscape and future prospects of antiviral drugs derived from microbial products. The Journal of Antibiotics 71, 45–52 (2018).

80 Zhang, B. et al. Crystal Structures of Membrane Transporter MmpL3, an Anti-TB Drug Target. Cell 176, 636–648.e613 (2019).

81 Andries, K. et al. A Diarylquinoline Drug Active on the ATP Synthase of Mycobacterium tuberculosis. Science 307, 223–227 (2005).

82 Zhang, Y. et al. Inhibition of M. tuberculosis and human ATP synthase by BDQ and TBAJ-587. Nature 631, 409–414 (2024).

83 Li, Q. et al. Functional and Structural Analysis of Influenza Virus Neuraminidase N3 Offers Further Insight into the Mechanisms of Oseltamivir Resistance. Journal of Virology 87, 10016–10024 (2013).

84 Alexander, J. A. N. et al. Structural and kinetic analyses of penicillin-binding protein 4 (PBP4)-mediated antibiotic resistance in Staphylococcus aureus. Journal of Biological Chemistry 293, 19854–19865 (2018).

85 Jeong, J.-H., Cha Hyung, J. & Kim, Y.-G. Crystal Structures of Penicillin-Binding Protein D2 from Listeria monocytogenes and Structural Basis for Antibiotic Specificity. Antimicrobial Agents and Chemotherapy 62, 10.1128/aac.00796–00718 (2018).

86 Cai, Y. et al. Distinct conformational states of SARS-CoV-2 spike protein. Science 369, 1586–1592 (2020).

87 Gobeil, S. M. C. et al. Effect of natural mutations of SARS-CoV-2 on spike structure, conformation, and antigenicity. Science 373, eabi6226 (2021).

88 Walls, A. C. et al. Structure, Function, and Antigenicity of the SARS-CoV-2 Spike Glycoprotein. Cell 181, 281–292.e286 (2020).

89 Xue, M. et al. Stable Attenuation of Human Respiratory Syncytial Virus for Live Vaccines by Deletion and Insertion of Amino Acids in the Hinge Region between the mRNA Capping and Methyltransferase Domains of the Large Polymerase Protein. Journal of Virology 94, 10.1128/jvi.01831–01820 (2020).

90 Turoňová, B. et al. In situ structural analysis of SARS-CoV-2 spike reveals flexibility mediated by three hinges. Science 370, 203–208 (2020).

91 Degen, D. et al. Transcription inhibition by the depsipeptide antibiotic salinamide A. eLife 3, e02451 (2014).

92 Ma, C., Yang, X. & Lewis Peter, J. Bacterial Transcription as a Target for Antibacterial Drug Development. Microbiology and Molecular Biology Reviews 80, 139–160 (2016).

93 Hibbs, R. E. & Gouaux, E. Principles of activation and permeation in an anion-selective Cys-loop receptor. Nature 474, 54–60 (2011).

94 Boyaci, H. et al. Fidaxomicin jams Mycobacterium tuberculosis RNA polymerase motions needed for initiation via RbpA contacts. eLife 7, e34823 (2018).

95 Szymański, M., Chmielewska, S., Czyżewska, U., Malinowska, M. & Tylicki, A. Echinocandins – structure, mechanism of action and use in antifungal therapy. Journal of Enzyme Inhibition and Medicinal Chemistry 37, 876–894 (2022).

96 Huang, X., Chen, M., Chen, Z. & Zhang, Y. Fungal β-1,3-glucan synthase: a review of structure, mechanism, and regulation. FEMS Yeast Research 25, foaf071 (2025).

97 Guo, H. et al. Structure of mycobacterial ATP synthase bound to the tuberculosis drug bedaquiline. Nature 589, 143–147 (2021).

98 Joon, S. et al. The NMR solution structure of Mycobacterium tuberculosis F-ATP synthase subunit ε provides new insight into energy coupling inside the rotary engine. FEBS Journal 285, 1111–1128 (2018).

99 Tan, Q. et al. Structure of the CCR5 Chemokine Receptor–HIV Entry Inhibitor Maraviroc Complex. Science 341, 1387–1390 (2013).

100 Lin, W. et al. Structural Basis of Transcription Inhibition by Fidaxomicin (Lipiarmycin A3). Molecular Cell 70, 60–71.e15 (2018).

101 Tupin, A., Gualtieri, M., Leonetti, J. P. & Brodolin, K. The transcription inhibitor lipiarmycin blocks DNA fitting into the RNA polymerase catalytic site. The EMBO Journal 29, 2527–2537 (2010).

102 Artsimovitch, I. et al. Allosteric Modulation of the RNA Polymerase Catalytic Reaction Is an Essential Component of Transcription Control by Rifamycins. Cell 122, 351–363 (2005).

103 Feklistov, A. et al. Rifamycins do not function by allosteric modulation of binding of Mg2+ to the RNA polymerase active center. Proceedings of the National Academy of Sciences 105, 14820–14825 (2008).

104 Sun, Q. et al. The molecular basis of pyrazinamide activity on Mycobacterium tuberculosis PanD. Nature Communications 11, 339 (2020).

105 Luckner, S. R., Liu, N., am Ende, C. W., Tonge, P. J. & Kisker, C. A Slow, Tight Binding Inhibitor of InhA, the Enoyl-Acyl Carrier Protein Reductase from Mycobacterium tuberculosis*. Journal of Biological Chemistry 285, 14330–14337 (2010).

106 Fisher, M. C. et al. Tackling the emerging threat of antifungal resistance to human health. Nature Reviews Microbiology 20, 557–571 (2022).

107 de Oliveira, H. C., Bezerra, B. T. & Rodrigues, M. L. Antifungal Development and the Urgency of Minimizing the Impact of Fungal Diseases on Public Health. ACS Bio & Med Chem Au 3, 137–146 (2023).

108 Fisher Matthew, C., et al. Threats Posed by the Fungal Kingdom to Humans, Wildlife, and Agriculture. mBio 11, 10.1128/mbio.00449–00420 (2020).

109 Nahirney, W. K. O. a. P. C. Netter’s Essential Histology: With Correlated Histopathology. 3rd edn, (Elsevier, 2020).

110 Abdella, S. et al. pH and its applications in targeted drug delivery. Drug Discovery Today 28, 103414 (2023).

111 Kim, N. pH variation impacts molecular pathways associated with somatic cell reprogramming and differentiation of pluripotent stem cells. Reproductive Medicine and Biology 20, 20–26 (2021).

112. Farley, A. R. & Link, A. J. in Methods in Enzymology Vol. 463 (eds Richard R. Burgess & Murray P. Deutscher) 725–763 (Academic Press, 2009).

113 Raju, M., Kavarthapu, R., Anbazhagan, R., Hassan, S. A. & Dufau, M. L. Blockade of GRTH/DDX25 Phosphorylation by Cyclic Peptides Provides an Avenue for Developing a Nonhormonal Male Contraceptive. Journal of Medicinal Chemistry 64, 14715–14727 (2021).

114 Zhong, Q., et al. Protein posttranslational modifications in health and diseases: Functions, regulatory mechanisms, and therapeutic implications. MedComm 4, e261 (2023).

115 Ramazi, S. & Zahiri, J. Post-translational modifications in proteins: resources, tools and prediction methods. Database 2021, baab012 (2021).

116 Suskiewicz, M. J. The logic of protein post-translational modifications (PTMs): Chemistry, mechanisms and evolution of protein regulation through covalent attachments. BioEssays 46, 2300178 (2024).

117 Singhal, A., Li, B. T. & O’Reilly, E. M. Targeting KRAS in cancer. Nature Medicine 30, 969–983 (2024).

118 Warren, H. R., Ross, S. J., Smith, P. D., Coulson, J. M. & Prior, I. A. Combinatorial approaches for mitigating resistance to KRAS-targeted therapies. Biochemical Journal 479, 1985–1997 (2022).

119 Perurena, N., Situ, L. & Cichowski, K. Combinatorial strategies to target RAS-driven cancers. Nature Reviews Cancer 24, 316–337 (2024).

120 Cregg, J. et al. Discovery of Daraxonrasib (RMC-6236), a Potent and Orally Bioavailable RAS(ON) Multi-selective, Noncovalent Tri-complex Inhibitor for the Treatment of Patients with Multiple RAS-Addicted Cancers. Journal of Medicinal Chemistry 68, 6064–6083 (2025).

