## Supplementary material for "Minimal Computational Framework for Systematic Identification of Antimicrobial Targets": The supplemental file includes a first page serving as an index describing its contents

Sergio A. Hassan
Bioinformatics and Computational Biosciences Branch, National Institutes of Allergy and Infectious Diseases, National Institutes of Health, U.S. DHHS, Bethesda, MD 20892

#### Overview

`pp.1-4:` Flowchart illustrating the pipeline architecture and overall workflow.
`pp.5-6:` General command syntax and instructions for session submission.
`pp.7-11:` Example commands and associated input data for the applications in the text.
`pp.12-13:` Example commands for all queries in the current implementation.
`pp.14-22:` Supplemental figures.

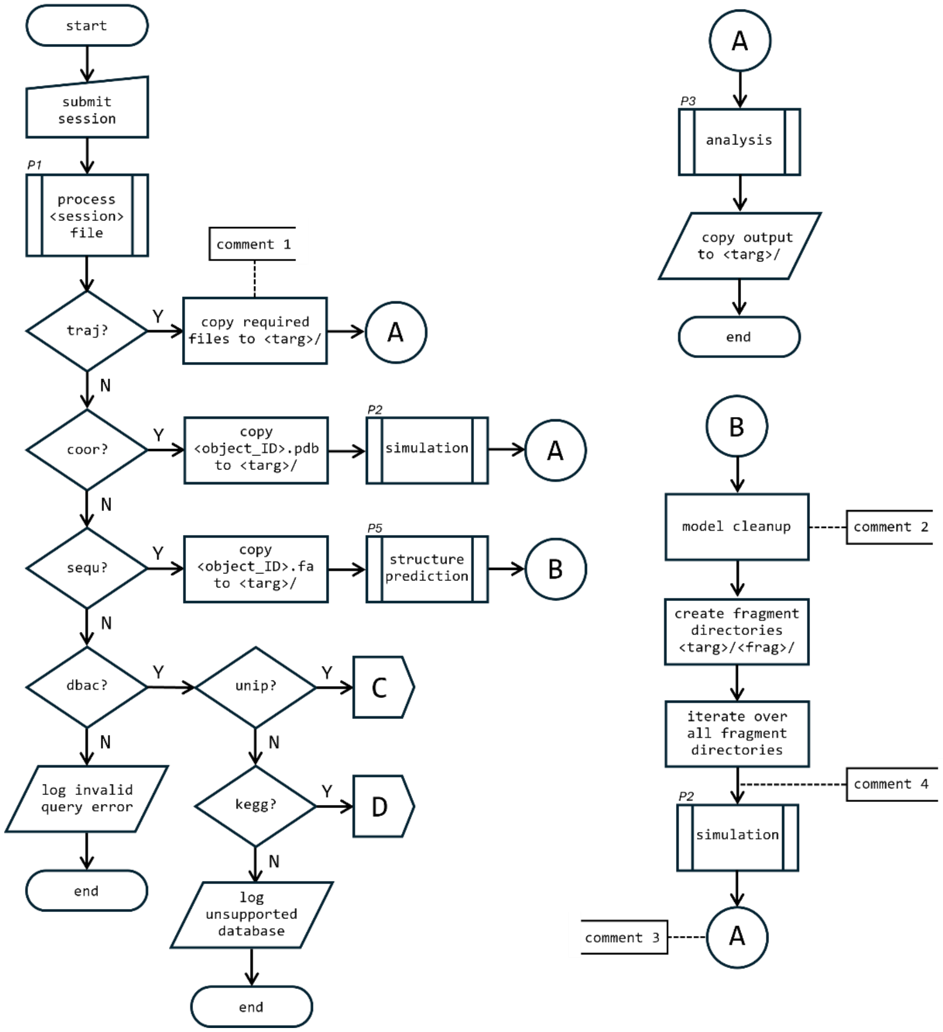
start
submit session
P1
process <session> file
traj?
Y
copy required files to <targ>/
comment 1
A
N
coor?
Y
copy <object_ID>.pdb to <targ>/
P2
simulation
A
N
sequ?
Y
copy <object_ID>.fa to <targ>/
P5
structure prediction
B
N
dbac?
Y
unip?
Y
C
N
kegg?
Y
D
N
log unsupported database
end
N
log invalid query error
end
A
P3
analysis
copy output to <targ>/
end
B
model cleanup
comment 2
create fragment directories <targ>/<frag>/
iterate over all fragment directories
comment 4
P2
simulation
comment 3
A

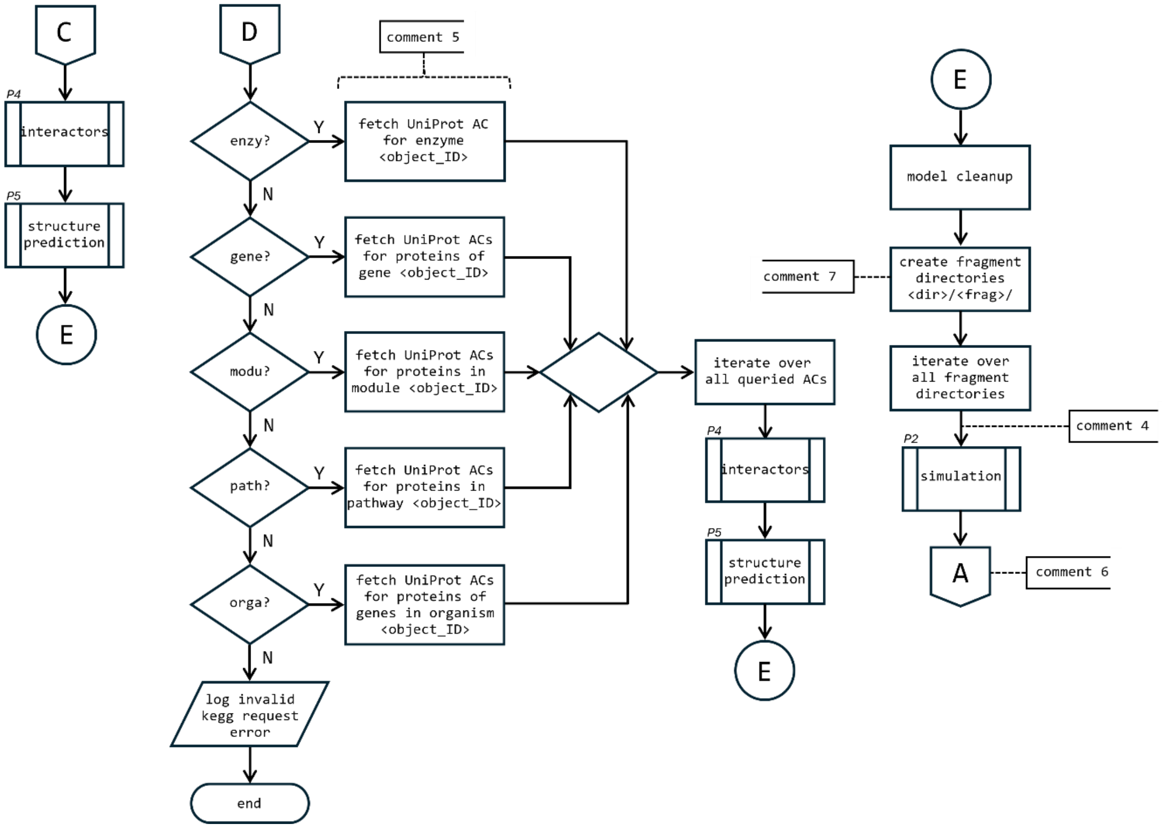
C
P4
interactors
P5
structure prediction
E
D
comment 5
enzy?
Y
fetch UniProt AC for enzyme <object_ID>
N
gene?
fetch UniProt ACs for proteins of gene <object_ID>
modu?
fetch UniProt ACs for proteins in module <object_ID>
path?
fetch UniProt ACs for proteins in pathway <object_ID>
orga?
fetch UniProt ACs for proteins of genes in organism <object_ID>
log invalid kegg request error
end
iterate over all queried ACs
P4
interactors
P5
structure prediction
E
E
model cleanup
comment 7
create fragment directories <dir>/<frag>/
iterate over all fragment directories
comment 4
P2
simulation
A
comment 6

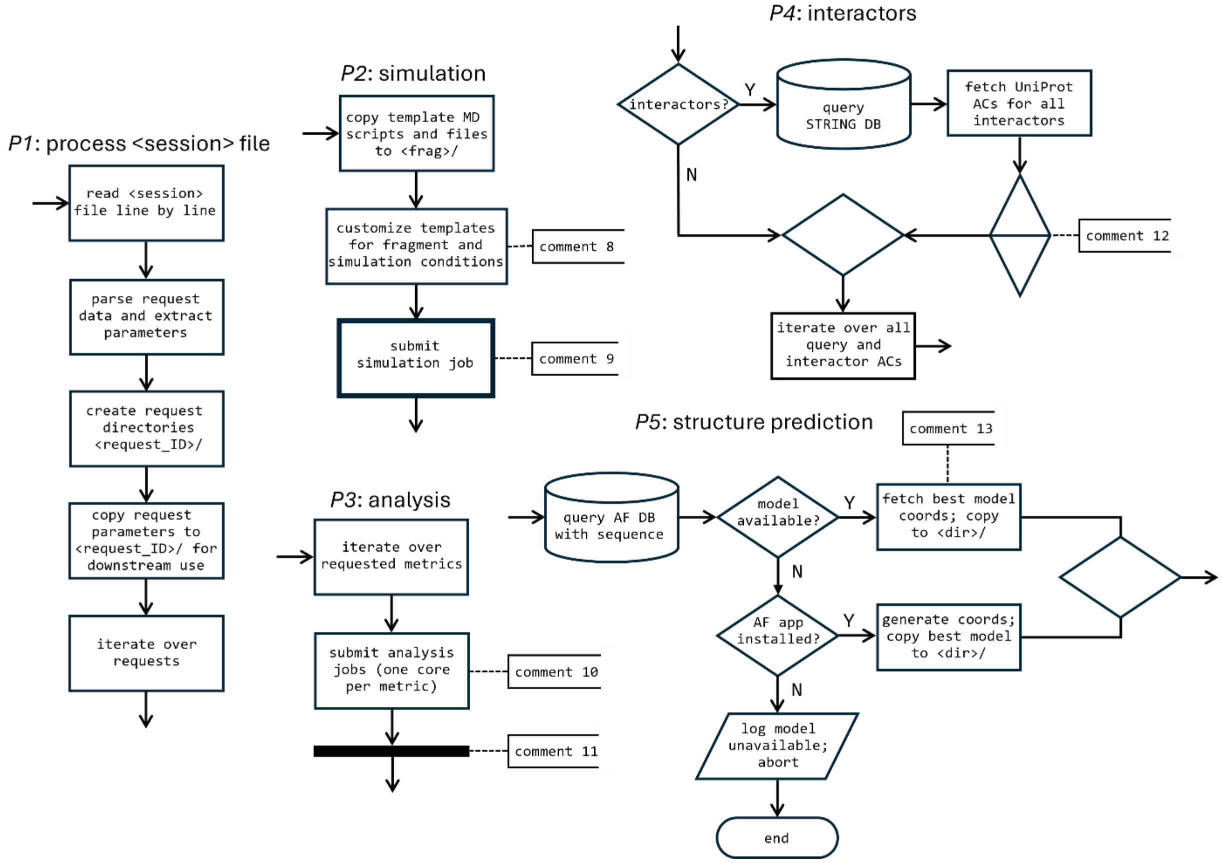
P1: process <session> file
read <session> file line by line
parse request data and extract parameters
create request directories <request_ID>/
copy request parameters to <request_ID>/ for downstream use
iterate over requests
P2: simulation
copy template MD scripts and files to <frag>/
customize templates for fragment and simulation conditions
comment 8
submit simulation job
comment 9
P3: analysis
iterate over requested metrics
submit analysis jobs (one core per metric)
comment 10
comment 11
P4: interactors
interactors?
Y
query STRING DB
fetch UniProt ACs for all interactors
N
comment 12
iterate over all query and interactor ACs
P5: structure prediction
comment 13
query AF DB with sequence
model available?
Y
fetch best model coords; copy to <dir>/
N
AF app installed?
Y
generate coords; copy best model to <dir>/
N
log model unavailable; abort
end

### Comments



comment 1: For query types traj, coor, and sequ: <targ>/ = <request_ID>/<object_ID>/
comment 2: Sanitization criteria described in main text.
comment 3: <targ>/ → <targ>/<frag>/
comment 4: Optional fragmentation step (e.g., PTM or pKa shifts; default: none; see main text).
If branching occurs, <frag>/ applies to each resulting subdirectory.
comment 5: ACs retrieved through suitable API queries to the KEGG DB.
comment 6: Same as flow A in page 1 but with <targ>/ → <dir>/<frag>/
comment 7: If query type = sequ: <dir>/ = <request_id>/<object_id>/
If query type = kegg & object type = orga: <dir>/ = <request_id>/<object_id>/<gene>/<UniProt_AC>/
If query type = kegg & object type ≠ orga: <dir>/ = <request_id>/<object_id>/<UniProt_AC>/
comment 8: Each <frag>/ is fully self-contained, with no external dependencies, and can be decoupled as an independent simulation directory.
System customization includes optimization of simulation cell size, and fragment and solvent preparation (see main text).
comment 9: Runs on multi-node HPC (high-speed interconnect) or multi-core single-node in HTC.
Most time-consuming step (hours to days depending on system size).
comment 10: Default mode: serial execution to support HPC and HTC resources.
For large systems, each metric is naïvely parallelized across multiple cores.
comment 11: Synchronization: wait until all metrics complete. Single incoming line represents multiple parallel metric flows.
comment 12: Sort ACs by node importance in the network (default: degree; other centrality metrics allowed; see main text).
comment 13: See definition of <dir>/ in comment 5.

The term “target” may refer to a genus (e.g., *Candida* vs. *Escherichia*), species (e.g., *C. albicans* vs. *C. auris*), biological network (e.g., metabolic or signal transduction), functional subnetwork or specific pathway (e.g., lipid or amino acid biosynthesis), network hub or edge, protein, protein binding site (e.g., orthosteric or allosteric), or any of its conformational substates (conformers). The workflow and associated commands enable analysis across all these target definitions. The method excludes membranes, an important class of targets for both known and emerging antimicrobials, e.g. antimicrobial peptides (AMPs), because they require a distinct treatment (see main text).

The core of the workflow comprises the NAMD molecular dynamics engine[1] and the CHARMM force field.[2] The implementation relies on CHARMM-compatible output formats. The underlying simulation platforms, including NAMD and CHARMM for classical simulations and GAMESS[3]–CHARMM for QM/MM (the latter representing an intended extension; see application to proton transfer reactions following analysis of the IWN metric[4]) are natively implemented in Fortran and C, making it desirable to maintain internal consistency for robustness and future integration. Consequently, the implementation consists of Fortran-based routines for data pre- and post-processing, with Linux (primarily Bash) scripting used for workflow orchestration; a directory containing equivalent implementations in C is also provided. Tasks involving interfacing with external databases and AI-based resources are more effectively handled using higher-level scripting languages commonly used in bioinformatics pipelines; however, this functionality is not implemented here, and the expander modules therefore remain in Fortran and Linux. For users unfamiliar with low-level compiled languages, an additional folder includes Python versions of the code generated using ChatGPT, provided for convenience and not independently validated.

External data integration includes AlphaFold2[5] and its local deployment, as well as STRING,[6] KEGG,[7] and UniProt.[8] The workflow employs GNUplot (http://www.gnuplot.info/) for plotting and generates scripts for structural and metric visualization in ChimeraX,[9] as well as network visualization in Cytoscape.[10] All components have been implemented, tested, and described in the main text, this SI, and the documentation and examples directory of the prototype (DOI: 10.17605/OSF.IO/EM2WH). While the current implementation makes explicit use of these software packages and databases, these choices are intended as concrete instantiations rather than fixed dependencies. A central design principle of the framework is modularity and functional decoupling, ensuring that individual components can be replaced, extended, or updated without altering the overall architecture or logic of the pipeline (see README in the distribution), thereby enabling adaptation to alternative or emerging tools and resources.

Software for downstream applications includes molecular docking using AutoDock Vina,[11] as well as de novo peptide design using RFdiffusion[12] and ProteinMPNN.[13] These tools are provided for convenience and are not part of the conceptual framework of the workflow, which is focused on target identification rather than lead discovery or molecular design.

**Session requests**

Requests are initiated using a session file with the command $./TBP.sh <session>. This can be run from a terminal or submitted as a job through the system's batch scheduler, depending on the computing environment (HPC or HTC; see README in the prototype distribution). The session file is a plain-text document containing a series of single-line requests, each in the following form:

```
[request_id] QUERy [query_type] [object_type] ID [object_id]
             SIMUlate [simulation_type]
             CALCulate [calculation_type] {metrics}
             INTEractors [protein interactors]
             ISIMulate [interactor simulation_type]
             ICALculate [interactor calculation_type] {metrics}
             ! (comments)
```

[request_id]: user-defined alphanumeric ID; multiple allowed (one per line); creates dedicated directory request.[request_id]/

[query_type]:traj|coor|sequ|unip|kegg
traj: requires simulation files x.pdb, x.psf (structure), x.dcd (trajectory), x.rtf (topology), and x.prm (parameters)
coor: requires model coordinates x.pdb, where x =[object_id]
sequ: requires amino acid sequence x.fa in fasta format; monomers only
unip and kegg: trigger API queries to external databases

[object_type]:prot|fast|orga|path|modu|gene|enzy
prot: for query types coor, traj, and unip
fast: for sequ
orga, path, modu, gene, and enzy: for kegg

[object_id]:(x)|AC|IDs
(x): user-provided file name for query type coor, sequ, or traj
AC: UniProt accession code
IDs: KEGG identifiers for object type orga, path, modu, gene, enzy

[simulation_type]:none|md
none: bypasses simulations for all queried proteins (mandatory for traj); only TOP is calculated on fixed fragment conformations if selected
md: requests simulations for all queried proteins; modular design allows alternative ensemble methods if compatible with selected metrics

[calculation type]:metrics
only implemented option

{metrics}:none|all|{list}
none: indicates no metric calculations
all: processes all metrics
{list}: single-line, unordered subset of metrics specified by 3-letter codes

[protein interactors]:none|all
none: no interactors processed; applies only to dbac
all: fetches interactors from the STRING database

[interactor simulation_type]:none|md

```
[interactor calculation_type]:metrics
{interactor metrics}:none|all|{list}
```

As in the QUERy parameters. Applies to all interactors if INTE=all

!(comments): unformatted user text; unlimited length

**Applications commands** (used in the main text)

**III.1:** Session submitted with:

```
$ ./TBP.sh HRV-B.str > HRV-B.log
```

where HRV-B.str contains:

```
HRVB3 QUER SEQU PROT ID hrv_b3 SIMU MD CALC METR ALL INTE NONE ISIM NONE ICAL IMET NONE ! human rhinovirus B serotype 3 (UniProt AC Q82081)

HRVB14 QUER SEQU PROT ID hrv_b14 SIMU MD CALC METR ALL INTE NONE ISIM NONE ICAL IMET NONE ! human rhinovirus B serotype 14 (UniProt AC P03303)

HRVB72 QUER SEQU PROT ID hrv_b72 SIMU MD CALC METR ALL INTE NONE ISIM NONE ICAL IMET NONE ! human rhinovirus B serotype 14 (NCBI AC ARE30262.1)
```

indicating that no interactors will be processed, and all metrics will be calculated for the queried fragments. The FASTA sequences [object_id].fasta for each request (see below) must be placed in the current working directory ($PWD) before submitting the session. Upon submission, the directories request.HRVB3/, request.HRVB14/, and request.HRVB72/ are created under $PWD, with results stored in subdirectories following the structure <protein>/<fragment>/<results>.

Once a fragment is processed (wall times may vary), RCO of the selected pockets can be calculated inside the <results> directory for that fragment. For the application discussed in the text (frags. X and VP2):

```
$ ./get_hotspot_RCO.sh fragment_X.hsp
$ ./get_hotspot_RCO.sh fragment_VP2.hsp
```

where fragment_X.hsp contains:

```
56 12.0 ! large pocket of X
27 10.0 ! small pocket of X
```

and fragment_VP2.hsp contains:

```
228 15.0 ! VP2:VP3
 82 11.0 ! VP2:VP2
116 13.0 ! VP2:VP1
  5 15.0 ! Allosteric, probably against VP1 and VP4
```

Residue numbering follows that of the fragment, not the polyprotein.

**III.2:** Session submitted with:

```
$./TBP.sh Mtb.str > Mtb.log
```

where `Mtb.str` contains:

```
MycoAcid QUER KEGG PATH ID mtu:mtu00074 SIMU MD CALC METR ALL INTE ALL
ISIM NONE ICAL IMET NONE ! mycolic acid biosynthesis for mAGP

ArabGalac QUER KEGG PATH ID mtu:mtu00572 SIMU MD CALC METR ALL INTE ALL
ISIM NONE ICAL IMET NONE ! Arabinogalactan biosynthesis for mAGP
```

This setup indicates that interactors of all proteins in both networks will be queried, but simulations and metric calculations will be performed only for the proteins listed in the query.

**III.3**: Session submitted with:

```
$./TBP.sh S.aureus.str > S.aureus.log
```

where `S.aureus.str` contains:

```
mono1 QUER UNIP PROT ID Q5HEB7 SIMU MD CALC METR ALL INTE NONE ISIM NONE
ICAL IMET NONE ! replica 1 of monomer

mono2 QUER UNIP PROT ID Q5HEB7 SIMU MD CALC METR ALL INTE NONE ISIM NONE
ICAL IMET NONE ! replica 2 of monomer

mono3 QUER UNIP PROT ID Q5HEB7 SIMU MD CALC METR ALL INTE NONE ISIM NONE
ICAL IMET NONE ! replica 3 of monomer

DimeTraj QUER TRAJ PROT ID dimer SIMU NONE CALC METR ALL INTE NONE ISIM
NONE ICAL IMET NONE ! dynamic trajectory of dimer created previously
```

This session submits four requests: the first three correspond to the monomer (UniProt AC Q5HEB7) and generate three independent analyses (to be combined after completion), while the fourth request queries a trajectory of the dimer model created previously. All metrics are calculated, and no interactors are included. Results will be in the corresponding fragment subdirectories under request.mono*/ and request.DimeTraj/

After the analyses of the three monomer replicas are complete, their metrics are combined using:

```
$ ./combine.sh monomers-1-2.str
$ ./combine.sh monomers-1-2-3.str
```

where `monomers-1-2.str` and `monomers-1-2-3.str` instruct the pipeline to combine the metrics of requests `mono1` and `mono2`, and `mono1`, `mono2`, and `mono3`, respectively. These files contain:

```
mono1 mono2  > MONO12  ! two combinations
mono1 mono2 mono3 > MONO123  ! three combinations
```

creating the directories `combined.requests.MONO12/` and `combined.requests.MONO123/` under `$PWD`, where the combined metrics are stored.

hrv_b3.fasta:

```
>sp|Q82081|POLG_HRV3 Genome polyprotein OS=Human rhinovirus 3
MGAQVSTQKSGSHENQNILTNGSNQTFTVINYYKDAASSSSAGQSFSMDPSKFTEPVKDL
MLKGAPALNSPNVEACGYSDRVQQITLGNSTITTQEAANAIVCYAEWPEYLSDNDASDVN
KTSKPDISVCRFYTLDSKTWKATSKGWCWKLPDALKDMGVFGQNMFYHSLGRTGYTIHVQ
CNATKFHSGCLLVVVIPEHQLASHEGGTVSVKYKYTHPGDRGIDLDTVEVAGGPTSDAIY
NMDGTLLGNLLIFPHQFINLRTNNTATIVVPYINSVPIDSMTRHNNVSLMVVPIAPLNAP
TGSSPTLPVTVTIAPMCTEFTGIRSRSIVPQGLPTTTLPGSGQFLTTDDRQSPSALPSYE
PTPRIHIPGKVRNLLEIIQVGTLIPMNNTGTNDNVTNYLIPLHADRQNEQIFGTKLYIGD
GVFKTTLLGEIAQYYTHWSGSLRISLMYTGPALSSAKIILAYTPPGTRGPEDRKEAMLGT
HVVWDIGLQSTIVMTIPWTSGVQFRYTDPDTYTSAGYLSCWYQTSLILPPQTSGQVYLLS
FISACPDFKLRLMKDTQTISQTDALTEGLSDELEEVIVEKTKQTLASVSSGPKHTQSVPA
LTANETGATLPTRPSDNVETRTTYMHFNGSETDVESFLGRAACVHVTEIKNKNAAGLDNH
RKEGLFNDWKINLSSLVQLRKKLELFTYVRFDSEYTILATASQPEASSYSSNLTVQAMYV
PPGAPNPKEWDDYTWQSASNPSVFFKVGETSRFSVPFVGIASAYNCFYDGYSHDDPDTPY
GITVLNHMGSMAFRVVNEHDVHTTIVKIRVYHRAKHVEAWIPRAPRALPYVSIGRTNYPR
DSKTIIKKRTNIKTYGLGPRFGGVFTSNVKIINYHLMTPDDHLNLVAPYPNRDLAVVATG
AHGAETIPHCNCTSGVYYSRYYRKFYPIICERPTNIWIEGSSYYPSRYQAGVMKGVGPAE
PGDCGGILRCIHGPIGLLTAGGGGYVCFADIRQLDFIADEQGLGDYITSLGRAFGTGFTD
QISAKVCELQDVAKDFLTTKVLSKVVKMISALVIICRNHDDLVTVTATLALLGCDGSPWR
FLKMYISKHFQVPYIERQANDGWFRKFNDACNAAKGLEWIANKISKLIEWIKNKVLPQAR
EKLEFCSKLKQLDILERQIASIHDSNPTQEKREQLFNNVLWLEQMSQKFSPLYASEAKRI
RDLKNKITNYMQFKSKQRTEPVCVLIHGTPGSGKSLTTSIVGRALAEHFNSSVYSLPPDP
KHFDGYQQQEVVIMDDLNQNPDGQDISMFCQMVSSVDFLPPMASLDNKGMLFTSNFVLAS
TNSNTLSPPTILNPEALIRRFGFDLDICMHSTYTKNGKLNAAMATSLCKDCHQPSNFKKC
CPLVCGKAISLVDRVSNVRFSIDQLVTAIINDYKNKVKITDSLEVLFQGPVYKDLEIDIC
NTPPPECISDLLKSVDSEEVREYCKKKKWIIPQISTNIERAVNQASMIINTILMFVSTLG
IVYVIYKLFAQTQGPYSGNPVHNKLKPPTLKPVVVQGPNTEFALSLLRKNILTITTEKGE
FTSLGIHDRICVLPTHAQPGDNVLVNGQKIQIKDKYKLVDPDNTNLELTIIELDRNEKFR
DIRGFISEDLEGLDATLVVHSNGFTNTILDVGPITMAGLINLSNTPTTRMIRYDYPTKTG
QCGGVLCTTGKIFGIHVGGNGRRGFSAQLKKQYFVEKQGLIVSKQKVRDIGLNPINTPTK
TKLHPSVFYNVFPGSKQPAVLNDNDPRLEVKLAESLFSKYKGNVQMEPTENMLIAVDHYA
GQLMSLDISTKELTLKEALYGVDGLEPIDVTTSAGYPYVSLGIKKRDILNKETQDVEKMK
FYLDKYGIDLPLVTYIKDELRSVDKVRLGKSRLIEASSLNDSVNMRMKLGNLYKAFHQNP
GIITESAVGCDPDVFWSVIPCLMDGHLMAFDYSNFDASLSPVWFECLEKVLNKLGFKQPS
LIQSICNTHHIFRDEIYRVEGGMPSGCSGTSIFNSMINNIIIRTLILDAYKGIDLDSLRI
LAYGDDLIVSYPFELDSNILAAIGKNYGLTITPPDKSDAFTKITWENITFLKRYFRPDPQ
FPFLIHPVMPMQDIYESIRWTRDPRNTQDHVRSLCMLAWHSGEKDYNDFITKIRTTDIGK
CLNLPEYSVLRRRWLDLF
```

hrv_b14.fasta :

```
>sp|P03303|POLG_HRV14 Genome polyprotein OS=Human rhinovirus 14
MGAQVSTQKSGSHENQNILTNGSNQTFTVINYYKDAASTSSAGQSLSMDPSKFTEPVKDL
MLKGAPALNSPNVEACGYSDRVQQITLGNSTITTQEAANAVVCYAEWPEYLPDVDASDVN
KTSKPDTSVCRFYTLDSKTWTTGSKGWCWKLPDALKDMGVFGQNMFFHSLGRSGYTVHVQ
CNATKFHSGCLLVVVIPEHQLASHEGGNVSVKYTFTHPGERGIDLSSANEVGGPVKDVIY
NMNGTLLGNLLIFPHQFINLRTNNTATIVIPYINSVPIDSMTRHNNVSLMVIPIAPLTVP
TGATPSLPITVTIAPMCTEFSGIRSKSIVPQGLPTTTLPGSGQFLTTDDRQSPSALPNYE
PTPRIHIPGKVHNLLEIIQVDTLIPMNNTHTKDEVNSYLIPLNANRQNEQVFGTNLFIGD
```

GVFKTTLLGEIVQYYTHWSGSLRFSLMYTGPALSSAKLILAYTPPGARGPQDRREAMLGT
HVVWDIGLQSTIVMTIPWTSGVQFRYTDPDTYTSAGFLSCWYQTSLILPPETTGQVYLLS
FISACPDFKLRLMKDTQTISQTVALTEGLGDELEEVIVEKTKQTVASISSGPKHTQKVPI
LTANETGATMPVLPSDSIETRTTYMHFNGSETDVECFLGRAACVHVTEIQNKDATGIDNH
REAKLFNDWKINLSSLVQLRKKLELFTYVRFDSEYTILATASQPDSANYSSNLVVQAMYV
PPGAPNPKEWDDYTWQSASNPSVFFKVGDTSRFSVPYVGLASAYNCFYDGYSHDDAETQY
GITVLNHMGSMAFRIVNEHDEHKTLVKIRVYHRAKHVEAWIPRAPRALPYTSIGRTNYPK
NTEPVIKKRKGDIKSYGLGPRYGGIYTSNVKIMNYHLMTPEDHHNLIAPYPNRDLAIVST
GGHGAETIPHCNCTSGVYYSTYYRKYYPIICEKPTNIWIEGNPYYPSRFQAGVMKGVGPA
EPGDCGGILRCIHGPIGLLTAGGSGYVCFADIRQLECIAEEQGLSDYITGLGRAFGVGFT
DQISTKVTELQEVAKDFLTTKVLSKVVKMVSALVIICRNHDDLVTVTATLALLGCDGSPW
RFLKMYISKHFQVPYIERQANDGWFRKFNDACNAAKGLEWIANKISKLIEWIKNKVLPQA
KEKLEFCSKLKQLDILERQITTMHISNPTQEKREQLFNNVLWLEQMSQKFAPLYAVESKR
IRELKNKMVNYMQFKSKQRIEPVCVLIHGTPGSGKSLTTSIVGRAIAEHFNSAVYSLPPD
PKHFDGYQQQEVVIMDDLNQNPDGQDISMFCQMVSSVDFLPPMASLDNKGMLFTSNFVLA
STNSNTLSPPTILNPEALVRRFGFDLDICLHTTYTKNGKLNAGMSTKTCKDCHQPSNFKK
CCPLVCGKAISLVDRTTNIRYSVDQLVTAIISDFKSKMQITDSLETLFQGPVYKDLEIDV
CNTPPPECINDLLKSVDSEEIREYCKKKKWIIPEIPTNIERAMNQASMIINTILMFVSTL
GIVYVIYKLFAQTQGPYSGNPPHNKLKAPTLRPVVVQGPNTEFALSLLRKNIMTITTSKG
EFTGLGIHDRVCVIPTHAQPGDDVLVNGQKIRVKDKYKLVDPENINLELTVLTLDRNEKF
RDIRGFISEDLEGVDATLVVHSNNFTNTILEVGPVTMAGLINLSSTPTNRMIRYDYATKT
GQCGGVLCATGKIFGIHVGGNGRQGFSAQLKKQYFVEKQGQVIARHKVREFNINPVNTPT
KSKLHPSVFYDVFPGDKEPAVLSDNDPRLEVKLTESLFSKYKGNVNTEPTENMLVAVDHY
AGQLLSLDIPTSELTLKEALYGVDGLEPIDITTSAGFPYVSLGIKKRDILNKETQDTEKM
KFYLDKYGIDLPLVTYIKDELRSVDKVRLGKSRLIEASSLNDSVNMRMKLGNLYKAFHQN
PGVLTGSAVGCDPDVFWSVIPCLMDGHLMAFDYSNFDASLSPVWFVCLEKVLTKLGFAGS
SLIQSICNTHHIFRDEIYVVEGGMPSGCSGTSIFNSMINNIIIRTLILDAYKGIDLDKLK
ILAYGDDLIVSYPYELDPQVLATLGKNYGLTITPPDKSETFTKMTWENLTFLKRYFKPDQ
QFPFLVHPVMPMKDIHESIRWTKDPKNTQDHVRSLCMLAWHSGEKEYNEFIQKIRTTDIG
KCLILPEYSVLRRRWLDLF

hrv_b72.fasta:

>ARE30262.1 polyprotein [rhinovirus B72]
MGAQVSTQKSGSHENQNILTNGSNQTFTVINYYKDAASSSSAGQSLSMDPSKFTEPVKDL
MLKGAPALNSPNVEACGYSDRVQQITLGNSTITTQEAANAVVCYAEWPEYLPDKDASDVN
KTSKPDTSVCRFYTLDSKMWQKNSKGWCWKLPDALKDMGVFGQNMFFHSLGRSGYTIHVQ
CNATKFHSGCLLVVVIPEHQLASHEGGNVSVKYKLTHPGEGGIDLSTNVEADGPVKDPVY
NMNGTLLGNLLIFPHQFINLRTNNTATLVVPYINSVPIDSMTRHNNVSLMIIPIAPLIVP
TNASPTLPITVTIAPMCTEFTGIRSKTVVPQGLPTTTLPGSGQFLTTDDRQSPSALPNYE
PTPRIHIPGQVKNLLEIIQVDTLIPMNNTHNRDEVKSYLIPLQPNRQGEMVFGTKLFIGD
GVFKTTLLGEIVQYYTHWAGSLRFSLMYTGPALSSAKLILSYTPPGANGPTTRKEAMLGT
HVVWDIGLQSTIVMTIPWTSGVQFRYTDPDTYTSAGFLSCWYQTSLILPPDTPNEVYMLS
FISACPDFKLRLMKDTQTISQTVALTEGLNDELEEVIVEKTKQTLASVSSGPKHTQSVPT
LTANETGATMPTLPSDNVETRTTYMHFNGSETDIECFLGRAACVHVTEIENKNPAGIDNH
KVGKLFNDWKISLSSLVQLRKKLELFTYVRFDSEYTILATASQPDTANYSSNLVVQAMYV
PPGAPNPVEWDDYTWQSASNPSVFFKVGDTSRFSVPYVGLASAYNCFYDGYSHDDANTPY
GISVLNHMGSMAFRIVNEHDTHRTLVKIRVYHRAKHIEAWIPRAPRALPYTSIGRTNYPK
NSQPVIKKRDGDIKTYGLGPRFGGVYTSNVKIMNYHLMTPEDHLNLVTPYPSRDLAVVAT
GAHGAEIIPHCNCTSGVYYSSYYRKFYPVICEKPTTIWIEGGPYYPSRFQVGVMKGVGPA
EPGDCGGILRCIHGPIGLLTAGGSGYVCFADIRQLECIAEEQGISDYITSLGRAFGVGFT
DQISAKVTELQDVAKDFLTTKVLSKVVKMVSALVIICRNYEDLVTVTATLALLGCDGSPW
RFLKMYISKHFQVPYIERQANDGWFRKFNDACNAAKGLEWIANKISKLIEWVKNKVLPQA
KEKLEFCSKLKQLDLLERQINTMHVSNPTQEKREQLFNNVLWLEQMSQKFAPLYATEAKR
IRELKNKIVNYMQFRNKQRTEPVCVLIHGTPGSGKSLTTSIIGRAIAEQFNSAVYSLPPD
PKHFDGYQQQEVVIMDDLNQNPDGQDISMFCQMVSSVDFLPPMASLDNKGMLFTSNFVLA
STNSNTLSPPTILNPDALARRFGFDLDICLHSTYTKNGKLNAAMATRVCKDCHQPTNFKK

```
CCPLVCGKAISLVDRATNVRYSIDQLVTAVVNDYKNKMQITDSLEVLFQGPVYKDLEIDV
NSTPPPECINDLLKSVDSEEIREYCKKKNWIIPEIPTNIERAVNQASMIINTILMFVSTL
GIIYVIYKLFAQTQGPYSGNPAHSKLKAPTLRPVVVQGPNTEFALSLLRKNIVTITTDKG
EFTGLGIYDHICVVPTHAQPNDNVLINGQKVQIKDKFKLVDPDNINLELTILDLNRNEKF
RDIRAFISEDFEGVEATLVVHSNNYTNTILEVGPVTMAGLINLSNTPTNRMIRYEYATKT
GQCGGVLCATGRIFGIHVGGNGRQGFSAQLKKQYFVEKQGQVIARQKVRELNINPVNTPT
KSKLHPSVFYNVFPGDKEPAVLSDNDPRLEVKLTESLFSKYKGNVEMEPTENMLVAVDHY
AGQLLSLDIPTEELTIKEALYGVDGLEPIDVTTSAGYPYVSLGIKKRDIINKETKDVEKM
KYYLDKYGIDLPLVTYIKDELRSTDKVRLGKSRLIEASSLNDSVNMRMKLGNLYKAFHQN
PGTITGSAVGCDPDVFWSVIPCLMDGHLMAFDYSNFDASLSPVWFVCLEKVLYKLGFKHS
SLIQSICNTHHIFRDEIYVVEGGMPSGCSGTSIFNSMINNIIIRTLVLDAYKGIDLDKLK
IIAYGDDLIVSYPYELNPEVLATLGKSYGLTITPPDKSATFSKITWENLTFLKRCFRPDK
QFPFLVHPVMPMKDIHESIRWTKDPKNTQDHVRSLCMLAWHSGEEEYNEFIGKIRTTDIG
KCLILPEYSVLRKRWLDLF
```

**General query types** (used in `examples/` in the prototype distribution)

```
coordie QUER COOR PROT ID j SIMU MD CALC METR TOP LME IWN DCN INTE NONE ISIM
NONE ICAL IMET NONE ! Model from PDB

1.prot QUER TRAJ PROT id mono SIMU none CALC METR PIN NIN INte none ISIM none
ICAL IMET none ! globin alpha

1.prot.1.lig QUER TRAJ PROT id monolig SIMU none CALC METR PIN NIN INte none
ISIM none ICAL IMET none ! globin alpha bound to HEME, Fe, and O2

4.prot QUER TRAJ PROT id tetra SIMU none CALC METR all INte none ISIM none
ICAL IMET none ! hemoglobin (tetramer; alpha/beta chains)

4.prot.4.lig QUER TRAJ PROT id tetralig SIMU none CALC METR all INte none
ISIM none ICAL IMET none ! oxygenated hemoglobin (tetramer + HEME + Fe + O2)

SEQUIX QUER SEQU FAST ID whateva SIMU MD CALC METR all INTE none ISIM NONE
ICAL IMET None ! sequence from GenBank

UPdb QUER UNIP PROT ID P02768 SIMU MD CALC METR ALL INTE all ISIM MD ICAL
IMET all ! asks uniprot

enzIE QUER kegg ENZY ID hsa:hsa00010:3.1.3.80 SIMU MD CALC METR all INTE ALL
ISIM MD ICAL IMET alL ! specific enzyme in pathway

genie QUER KEGG GEne ID hsa:131 SIMU MD CALC METR ALL INTE All ISIM Md ICAL
IMET all ! gene for human alcohol dehydrogenase

o0d_l QUER kegg MODU ID hsa:M00001 SIMU MD CALC METR top LME LCF PIN nin INTE
all ISIM NONE ICAL IMET NONE ! get module (subnetwork) in H. sapiens

path.y.Z QUER KEGG PATH id csyr:csyr00520 SIMU md CALC METR TOP LCF PIN INte
all ISIM none ICAL IMET none ! get sugar nucleotide metabolism pathway from a
primate (C. syrichta)

orGaneX QUER kegg ORGA ID mge SIMU none CALC METR NONE INTE none ISIM NONE
ICAL IMET NONE ! query smallest organism (all 476 genes will be processed =>
large MD batch submission)
```

**Design application**

Although the main goal of the method and its implementation is target identification, the following example illustrates an auxiliary feature in which a peptide is designed to target the two conformers ($c_1$ and $c_2$) of a hotspot in fragment VP2 (see Application III.1 and Fig. 5). It also demonstrates how interaction plots can be used to identify weak and strong interfacial contacts, with the aim of improving design.

Each conformer is treated as an independent target. The workflow provides scripts and templates for small molecule screening (AutoDock Vina) and de novo peptide design (RFdiffusion and ProteinMPNN). The example here focuses on cyclic peptides. Other design types can be implemented by modifying the templates generated by the workflow (see examples/ and doc.txt in the distribution). De novo design is carried out by submitting

```
$ ./optimizer.sh <denovo>
```

where <denovo> is a plain text input file (for example, design.str) with the single line form,

```
TARG [name] CENT [hspc] RADI [hspr] NUMP [bb] NUMS [ss] PEPL [nres] ! comment
```

Where:

[name]: basename of the target protein file ([name].pdb)
[hspc]: center of the sphere defining the hotspot conformer
[hspr]: radius (in Å) of that sphere
[nres]: peptide length (number of residues)
[bb]: number of backbone designs (poly-Gly) generated by RFdiffusion
[ss]: number of optimized sequences per backbone as generated by ProteinMPNN

Notes: [hspc] and [hspr] are typically the same values used in <hotspots> for determining the conformers (III.1); default values for [bb] and [ss] are 100 (i.e., 10,000 peptides are generated).

Using c1.pdb and c2.pdb for the main and secondary conformers of fragment VP2, the design_1.str and design_2.str files used here are

```
TARG c1 CENT 94 RADI 15.0 NUMP 20 NUMS 20 PEPL 50 ! comment
TARG c2 CENT 94 RADI 15.0 NUMP 20 NUMS 20 PEPL 50 ! comment
```

(see example/ and doc.txt in the distribution for details) The values 20 and 50 are chosen for illustration, but larger values are preferable. The pipeline proceeds through a series of stages: (1) RFdiffusion, (2) ProteinMPNN, (3) energy minimization, (4) interaction energy calculation ($E_{int}$), (5) ranking by $E_{int}$, (6) selection of top $E_{int}$, (7) molecular dynamics simulation, (8) RMSD versus time calculation, and (9) final selection for downstream analysis.

The workflow generates one dedicated directory for each optimized peptide, representing a subset of the 20 × 50 peptides as described below. Steps 1 and 2 use the two external software. Step 3 is performed with CHARMM on the all-atom complex (protein bound to the peptide) in vacuum. In step 4, $E_{int}$ is the protein–peptide interaction energy from step 4. Step 5 ranks $E_{int}$ from highest (most negative) to lowest. Step 6 selects the top binding modes, defined as all peptides with $E_{int}$ within the top $\Delta$ = 10 × [nres] kcal/mol, allowing 10 kcal/mol per residue in the peptide, a reasonable value to account for modulation due to water.[14] The selected complexes proceed to step 7, which is a full MD simulation in water as described in the main pipeline. The simulation is brief (2 ns) and is used to assess which binding modes are stable in the fully atomistic environment. In step 9, stability is evaluated using the peptide Cα-RMSD versus time. Stable peptides remain near their initial position, whereas unstable ones deviate. The Cα-RMSD is computed for each snapshot after Cα-superposition of the protein onto the reference protein [name].pdb. Only stable binding modes are considered for downstream analysis. This may include longer simulations and subsequent querying with `TRAJ` to obtain metrics for the complexes, including peptide–protein `PIN` and `NIN`, which the pipeline can generate as described below.

Design improvements can follow several directions. Given the large hotspot and relatively small peptides, optimal peptides can be fused to create polycyclic compounds, or extended with branches to better engage underused regions of the interface. Chemical modifications, such as incorporating unusual or unnatural amino acids, can also be introduced to expand the chemical space beyond the 20 natural amino acids while remaining within a manageable search space (see Fig. S9). These approaches are beyond the scope of the current method and workflow.

**References**


1 Phillips, J. C. *et al.* Scalable molecular dynamics on CPU and GPU architectures with NAMD. *The Journal of Chemical Physics* **153**, 044130 (2020).

2 Brooks, B. R. *et al.* CHARMM: The Biomolecular Simulation Program. *J. Comp. Chem.* **30**, 1545 (2009).

3 Barca, G. M. J. *et al.* Recent developments in the general atomic and molecular electronic structure system. *The Journal of Chemical Physics* **152**, 154102 (2020).

4 Lee, Y.-S., Hassan, S. A. & Hurt, D. E. Proton Transfer Mechanisms in the MmpL3 Transporter of Mycobacterium tuberculosis Studied by Computer Simulations. *ACS Bio & Med Chem Au* DOI: 10.1021/acsbiomedchemau.5c00274 (2026).

5 Jumper, J., Evans, R., Pritzel, A. & al., e. Highly accurate protein structure prediction with AlphaFold. *Nature* **596**, 583–589 (2021).

6 Szklarczyk, D. *et al.* The STRING database in 2025: protein networks with directionality of regulation. *Nucleic Acids Res.* **53**, D730–D737 (2025).

7 Kanehisa, M., Furumichi, M., Sato, Y., Matsuura, Y. & Ishiguro-Watanabe, M. KEGG: biological systems database as a model of the real world. *Nucleic Acids Res.* **53**, D672–D677 (2025).

8 The UniProt Consirtium, UniProt: the Universal Protein Knowledgebase in 2025. *Nucleic Acids Res.* **53**, D609–D617 (2025).

9 Pettersen, E. F. *et al.* UCSF ChimeraX: Structure visualization for researchers, educators, and developers. *Protein Science* **30**, 70–82 (2021).

10 Shannon, P. *et al.* Cytoscape: A Software Environment for Integrated Models of Biomolecular Interaction Networks. *Genome Research* **13**, 2498–2504 (2003).

11 Trott, O. & Olson, A. J. AutoDock Vina: Improving the speed and accuracy of docking with a new scoring function, efficient optimization, and multithreading. *Journal of Computational Chemistry* **31**, 455–461 (2010).

12 Watson, J. L. *et al.* De novo design of protein structure and function with RFdiffusion. *Nature* **620**, 1089–1100 (2023).

13 Dauparas, J. *et al.* Robust deep learning–based protein sequence design using ProteinMPNN. *Science* **378**, 49–56 (2022).

14 Hassan, S. A. Intermolecular Potentials of Mean Force of Amino Acid Side Chain Interactions in Aqueous Medium. *J. Phys. Chem. B* **108**, 19501–19509 (2004).

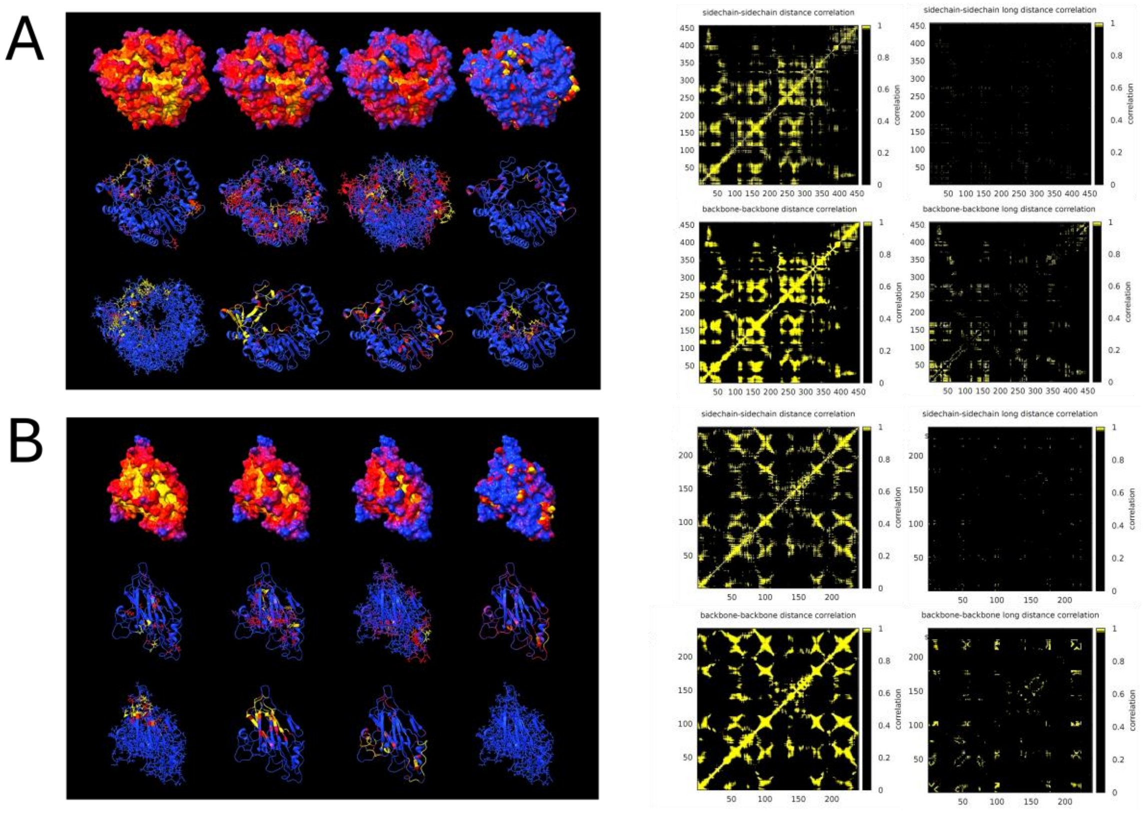


**Figure S1.** Metrics visualization. As in the upper panel of Figure 3A, shown for fragment 3D (A) and VP2 (B). Complete plots complementing the DCN metric mapped on the protein are shown on the right for each fragment, including $\texttt{DCN}_{\texttt{sc}}$, $\texttt{DCN}_{\texttt{sc,lr}}$, $\texttt{DCN}_{\texttt{bb}}$, and $\texttt{DCN}_{\texttt{bb,lr}}$ (index `lr` stands for long-range; see text and `doc.txt` in the distribution). As with all other metrics, detailed quantitative analysis and ligand design or screening can be performed using the plain text data files included in the output package (see Fig. S9 for a specific application).

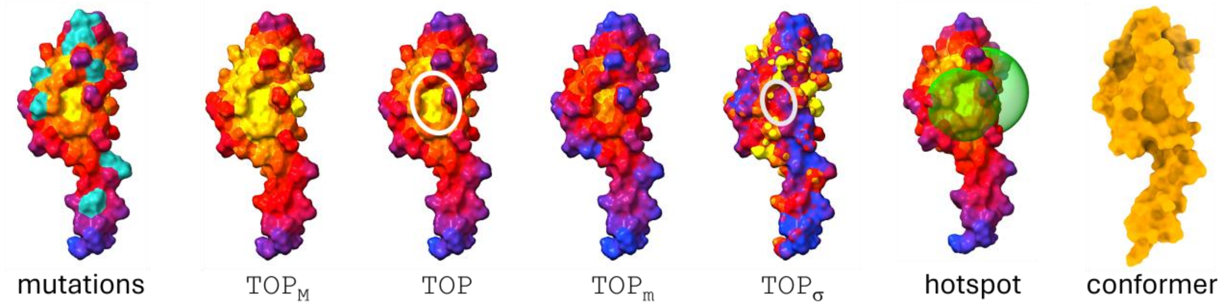


**Figure S2.** Same as Figure 4, for the second (smaller) crevice of fragment X. TOP analysis suggests the pocket undergoes breathing. The backbone conformation of the hotspot remains constant, as indicated by the single conformer (100% occupancy). Topography changes arise solely from side-chain movements, with LCF metrics indicating that these changes are sporadic and transient.

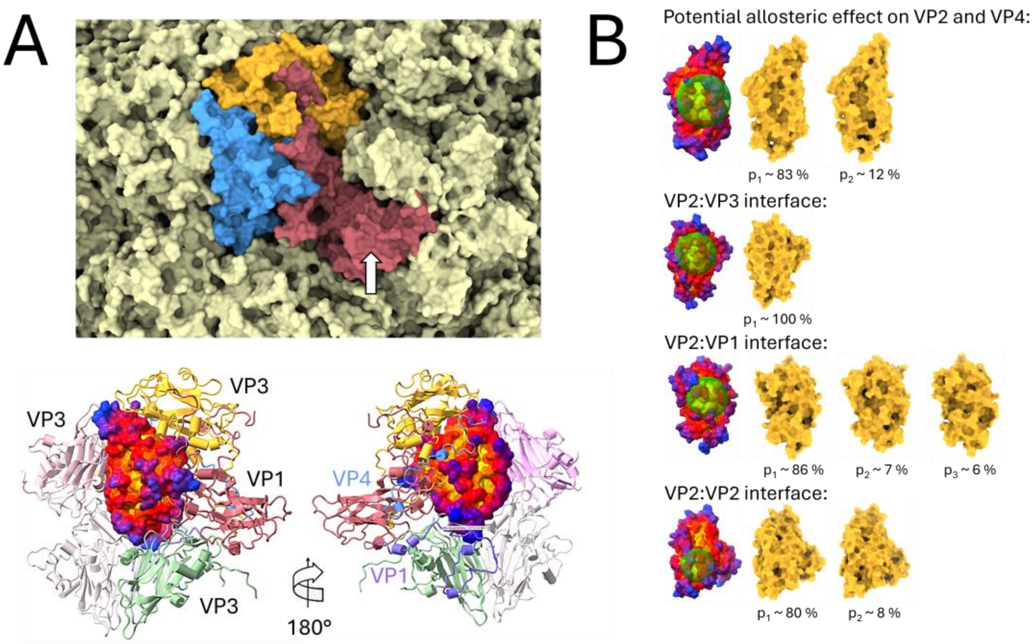


**Figure S3.** (A) Structural context of fragments VP2 (blue), VP3 (amber), and VP1 (rust) in the mature capsid of serotype B14, viewed from outside the virion (upper panel; PDB ID: 4RHV). The lower panel shows the `TOP` metric on VP2, highlighting proteins in direct contact with it in the mature virion (VP4 in blue); orientation of lower left panel same as in upper panel. (B) The four conserved pockets identified on VP2 from these metrics: one potential allosteric site (see Fig. 5 and an example of de novo design of a cyclic peptide in Fig. S9) and three interfacial sites with distinct pocket backbone conformations. Well-characterized antivirals (although none yet approved for therapy) bind in the interior of the outer-facing lobe of VP1, indicated by the arrow in (A) (e.g., PDB ID: 3VDD), increasing its rigidity and thereby hindering the conformational changes required for uncoating and RNA release during infection.

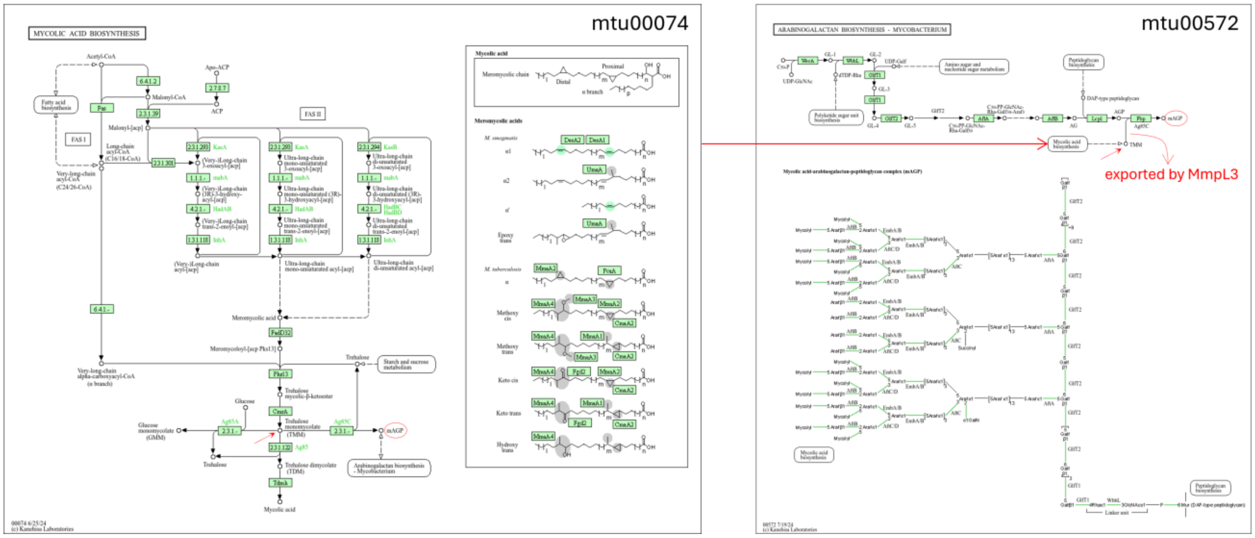


**Figure S4.** Schematic representations of the two KEGG pathways queried during pipeline execution that are involved in mAGP biosynthesis. The mAGP complex is shown in the lower panel of pathway mtu00572. Pathway mtu00074 is cytosolic and inner-membrane–associated, whereas mtu00572 comprises periplasm-facing reactions involved in mAGP assembly (note that Ag85C, although annotated in mtu00074, acts after TMM export by MmpL3). Labels and symbols shown in red were added for contextual clarity and were not generated by the pipeline: the horizontal arrow indicates the relationship between the pathways, inclined arrows represent the TMM molecule, and the oval denotes the final mAGP complex. The enzymes shown are localized to different cellular compartments, including the plasma membrane, the cytoplasm, and the cell-wall/periplasmic space. Diagrams are downloaded from the KEGG database by default during pipeline execution.

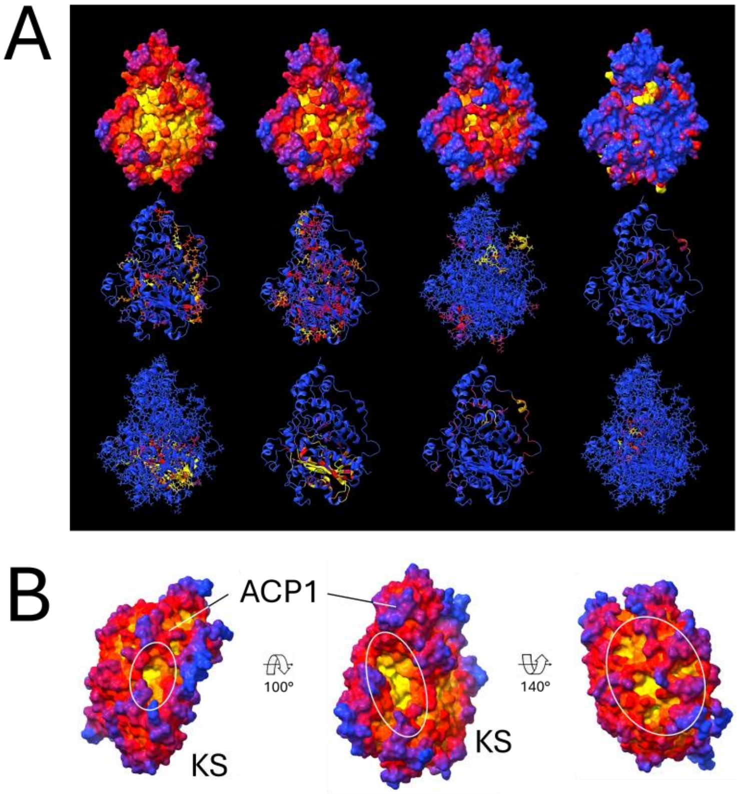


**Figure S5.** (A) Metrics for Fragment 1 (ACP1–KS) of the Pks13 protein analyzed in III.2. Pks13 is a polyketide synthase and corresponds to one of the central nodes of the mtu00074 mycolic acid biosynthesis network (Figs. 6 and S4; UniProt AC: I6X8D2). (B) Three hotspots identified using the metrics, with potentially distinct effects on network function upon ligand binding. The three regions (indicated by white ovals) represent three classes of hotspots of interest: a small pocket, a large deep crevice, and multiple closely spaced shallow pockets.

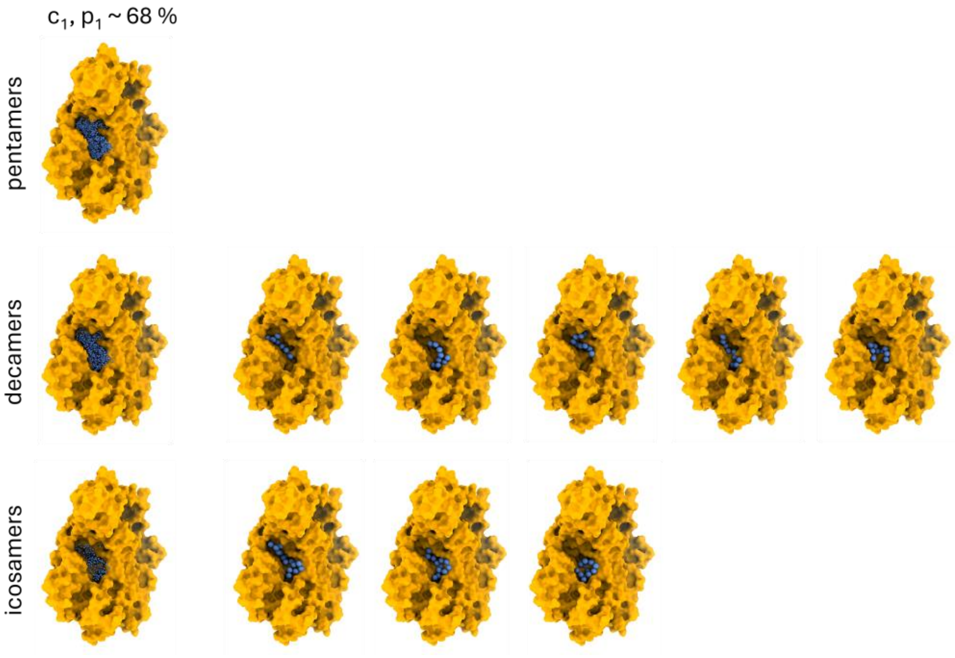


**Figure S6.** Internal water networks identified from the `IWN` metric mapped onto the main crevice (Fig. S6B, middle panel) of conformer 1 of fragment 1. The left column shows the aggregation of all water chains sampled over the dynamics for 5-mers, 10-mers, and 20-mers, and the panels at the right show representative chain species connecting polar groups within the pocket, some of which may be functionally relevant.[4] The workflow generates statistics on chain–fragment contacts (`IWN` metric) as well as network species and population distributions, and produces coordinates for both chains and protein structures for downstream analyses. Output data allows calculation of residence times.

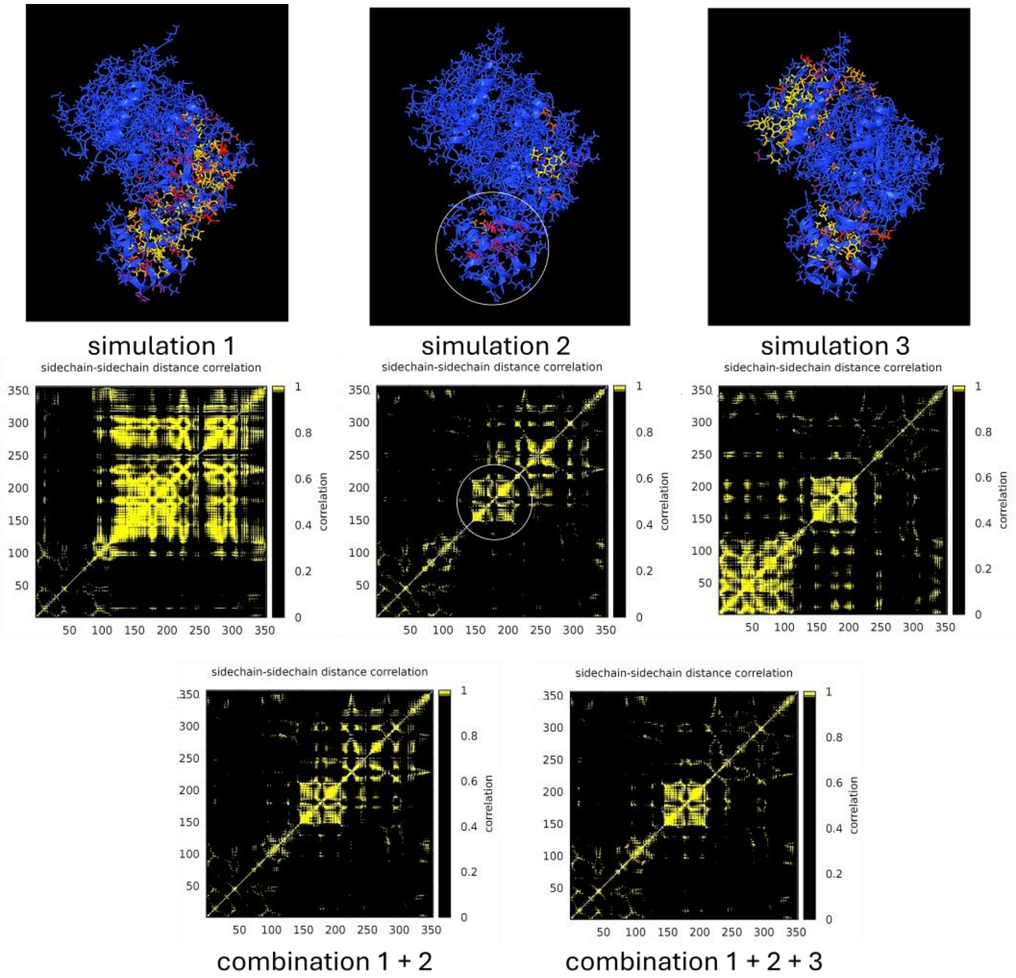


**Figure S7.** $\mathrm{DCN_{sc}}$ metric from three independent simulations of the monomer in application III.3. Cross-correlations are highly sensitive to simulation conditions, but appropriate combination of the data can mitigate statistical fluctuations and retain the common, physically meaningful correlations. Here, combining DCN matrices involves averaging normalized values for each residue pair (*i*,*j*) across simulations, so high correlations observed in only a few simulations are downweighted relative to smaller correlations consistently observed across most simulations. In this case, the smaller lobe (circled on the reference heatmap in the upper row; residues 145–215, corresponding to the central domain, CD) shows a common pattern repeated in all simulations (circled, middle row). In contrast, other regions become less defined as convergence across replicas increases, preserving the central pattern while diminishing high correlations observed in the N- and C-terminal domains (NTD and CTD, respectively). Other metrics are combined in a similar manner to yield statistically robust results.

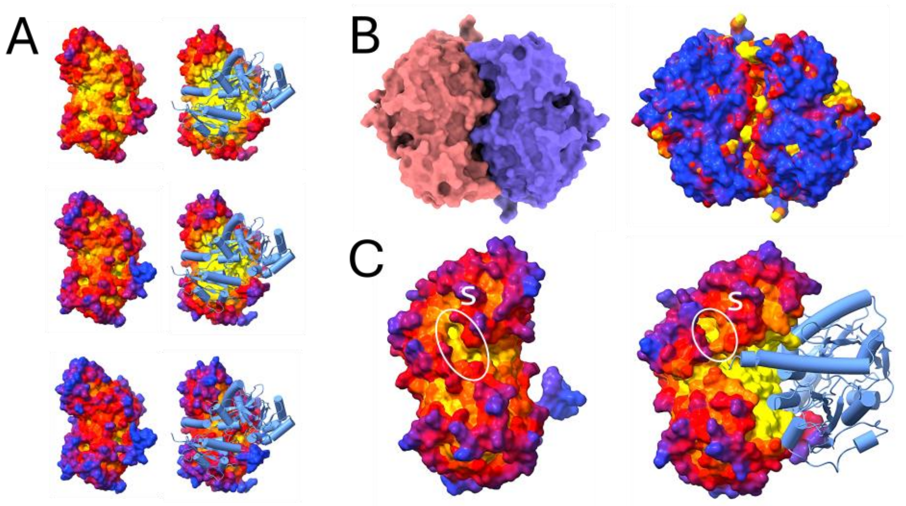


**Figure S8.** (A) Topography of Ddl as a free monomer and in the homodimeric state in application III.3: middle panel, TOP metric as in Fig. 7C; upper and lower panels show $TOP_M$ and $TOP_m$, respectively. (B) View of the dimer showing the two monomers (left) and $TOP_\sigma$, highlighting topographic fluctuations at the monomer–monomer interface; PIN and NIN (not shown) suggest that suitable ligands could perturb stabilizing interdimer interactions. (C) Potential hotspots (circled) not shown in Fig. 7D; these crevices may serve as entry pathways connecting to the ATP-binding site within the catalytic cleft.

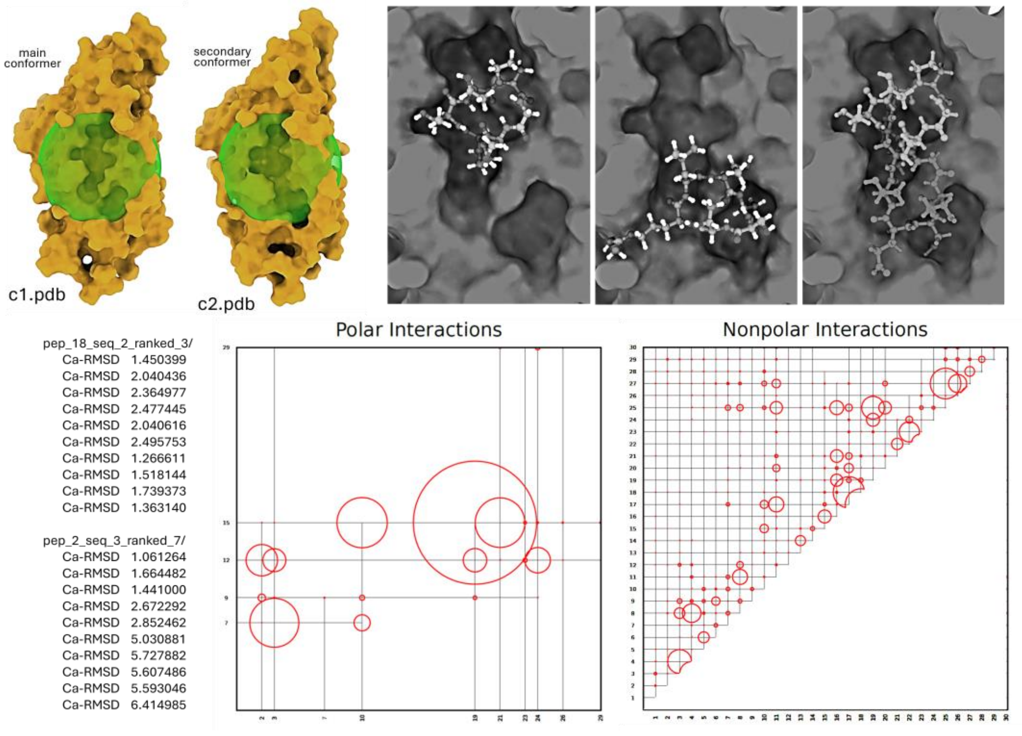


**Figure S9:** Downstream application illustrating auxiliary output scripts and features generated by the pipeline. The example shows de novo design of tail-to-head cyclic hexapeptides targeting the main and secondary conformers of a hotspot in structural protein VP2 in HRV-B14 (see Application III.1), with potential allosteric effects on VP2:VP4 and VP2:VP2 interfaces. Upper left: primary and secondary hotspot conformers with spheres defining their spatial extent, as determined by the `TOP` metric. Upper right: two representative hexapeptides targeting distinct regions of the hotspot, showing empty crevices that can be exploited by side-chain branching (not shown) or by combining high-avidity peptides to form polycyclic constructs, with a bicyclic peptide shown schematically for illustration. Lower left: Cα RMSD time series (2 ns) for two designed hexapeptides, one stable and one unstable; each peptide design is stored in a dedicated directory with naming convention `pep_[b]_seq_[s]_ranked_[r]/` where `[b]` = 1 ... `[bb]`, `[s]` = 1 ... `[ss]` (see `<denovo>` script), and `[r]` is the rank based on $E_{int}$, where lower `r` corresponds to larger $|E_{int}|$. Lower right: interaction-analysis outputs for `NIN` and `PIN` with circle radii proportional to interaction strength (see `doc.txt`); these plots can be used for comparative analysis (e.g., mutations, changes in solution conditions) and to guide peptide modification to enhance weaker interactions. The `NIN` plot is symmetric and represents residue-number vs residue-number in the complex; the `PIN` plot is asymmetric and represents acceptor residue-number vs donor residue-number. Plots are generated using `./plot_hb-interactions.x` or `./plot_hydpho-interactions.x` (the plots shown do not correspond to the protein–peptide system designed here and originate from a different application involving a miniprotein queried through `COOR`).
